# The choice of structural template can create apparent links between predicted protein stability and longevity in tetrapods

**DOI:** 10.64898/2026.10.12.758602

**Authors:** Enso Onill Torres Alegre, Diana E. Mora Jimenez

## Abstract

Comparative studies increasingly score amino-acid variation across species with structure-based predictors applied to a single structural template, but whether the template itself can create associations with species traits has received little attention. We examined this while testing whether long-lived species carry proteins predicted to be more stable or, under nearly neutral theory, less stable. In two independent tetrapod radiations, 57 placental mammals (1,755 genes) and 40 reptiles and amphibians (1,910 genes), we reconstructed ancestral proteins, predicted with ThermoMPNN the folding effect of every amino-acid difference between the ancestor and each species on AlphaFold backbones, and related the mean effect to lifespan by phylogenetic regression, following analysis plans fixed in advance. With a human backbone, primates, which are long-lived for their size, appeared least destabilized, producing a weak apparent association with lifespan. In a pre-specified confirmatory test, a platypus backbone reduced the primate offset from *−*1.96 to *−*0.62 SD and the association disappeared (*P* = 0.701). In reptiles, a human backbone, although equidistant in time, favoured slowly evolving turtles and crocodilians and gave a nominal association that two fish backbones did not reproduce. The predictor captured site-level stability signal, yet with outgroup templates neither radiation showed a robust association between predicted stability and longevity, although moderate effects cannot be excluded. The choice of structural template can therefore create apparent cross-species associations. We propose four checks to detect them: ancestral references, outgroup templates, tests for clade and rate effects, and tests of dependence on a few clades.

## 1. Introduction

Maximum lifespan varies about 100-fold across mammals, and ectothermic tetrapods span an even wider range of ageing rates, including phylogenetically widespread negligible senescence [1] and the slow ageing of turtles [2]. Among the molecular mechanisms proposed for this variation is the maintenance of protein stability and integrity. Naked mole-rat proteins resist urea-induced unfolding better than mouse proteins [3], SIRT6 variants of long-lived rodents repair DNA more efficiently [4], and a comparative analysis of mammalian genomes concluded that human longevity-associated proteins are more stable than their orthologs in short-lived mammals [5].

Molecular evolution offers a competing expectation. Large, long-lived species tend to have smaller effective population sizes, in which purifying selection is less efficient and slightly deleterious substitutions fix more often [6]; large mammals accumulate more radical substitutions in mitochondrial proteins [7]. If many slightly deleterious substitutions reduce folding stability, long-lived species could carry more destabilizing substitutions. This expectation depends on assumptions about effective population sizes and the distribution of substitution effects that we do not estimate here.

Testing these predictions requires a stability measure that varies meaningfully between orthologs. AlphaFold models of orthologs are nearly identical and largely insensitive to point substitutions [8, 9]. We therefore predicted the effect of each amino-acid difference on folding free energy with ThermoMPNN [10], trained on mega-scale stability measurements [11], and measured differences from reconstructed ancestors rather than from a modern reference species. Predictors of this kind score the variants of every species on one backbone, usually the structure of a model or reference species, and whether that choice can itself create associations with a species trait has received little attention. In the course of the first study we found unexpectedly that it can, and we then tested this bias with a confirmatory analysis and examined it again in a second radiation. The paper therefore addresses two questions: whether the predicted folding stability of amino-acid differences from an ancestor is associated with longevity, and how the structural template can bias such comparative analyses and how the bias can be detected. We ran the analysis in two independent radiations, each under its own analysis plan: placental mammals, which are endotherms, and reptiles and amphibians, which are ectotherms. Figure 1 summarizes the design and the main findings.

**Figure 1:**
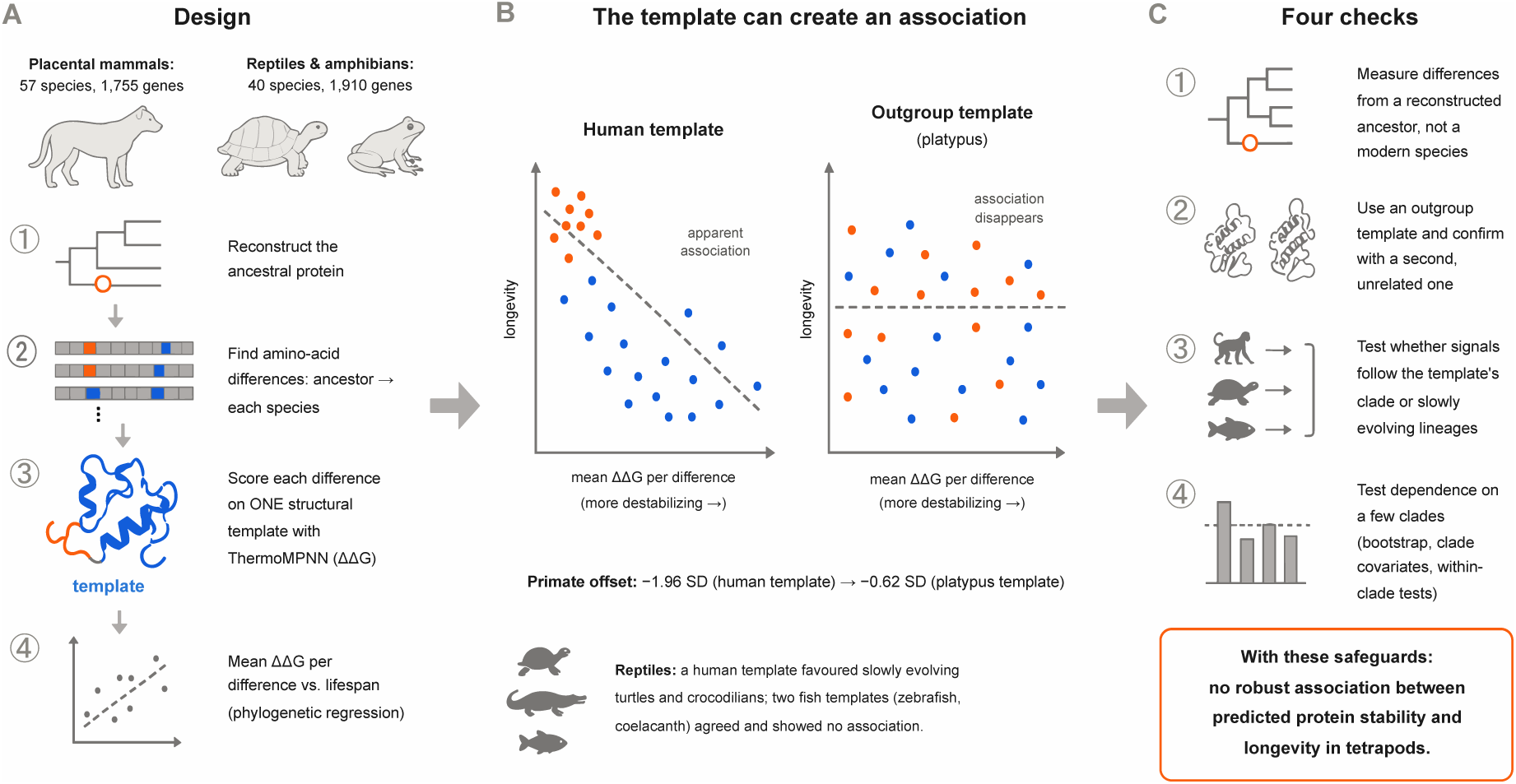
Overview of the study. (A) Design of the two comparative studies. (B) How the structural template can create an apparent association with longevity; the scatter plots are schematic and do not show data, whereas the primate offsets are those of the confirmatory test (Section 3.3). (C) The four checks proposed to detect this bias.

## 2. Methods

### 2.1 Design and pre-specified plans

Both studies followed the same logic (Table 1). For each species and gene, we identified the amino-acid differences between a reconstructed ancestor and the extant protein and predicted their effect on folding stability on a single structural backbone; species-level summaries were then related to lifespan. Each study had an internal analysis plan written before protein, ortholog or structure data were retrieved, and dated amendments were recorded. Plans were hashed with SHA256 at each stage; the hashes show that their contents were not altered afterwards but do not independently certify when they were written, and neither plan was deposited in a public registry. In the mammal study, three operational amendments preceded all association tests, one analysis (an opossum template) was added after the primary results were known and is labelled post hoc, and one confirmatory amendment (E5, a platypus template) was specified before any of its data were retrieved. In the reptile and amphibian study, a single operational amendment preceded all tests; its design incorporated the lessons of the mammal study. Software versions, data releases with retrieval dates, genome assemblies and all fixed parameters are given in the Supplementary Methods (Tables S1–S9).

**Table 1:**
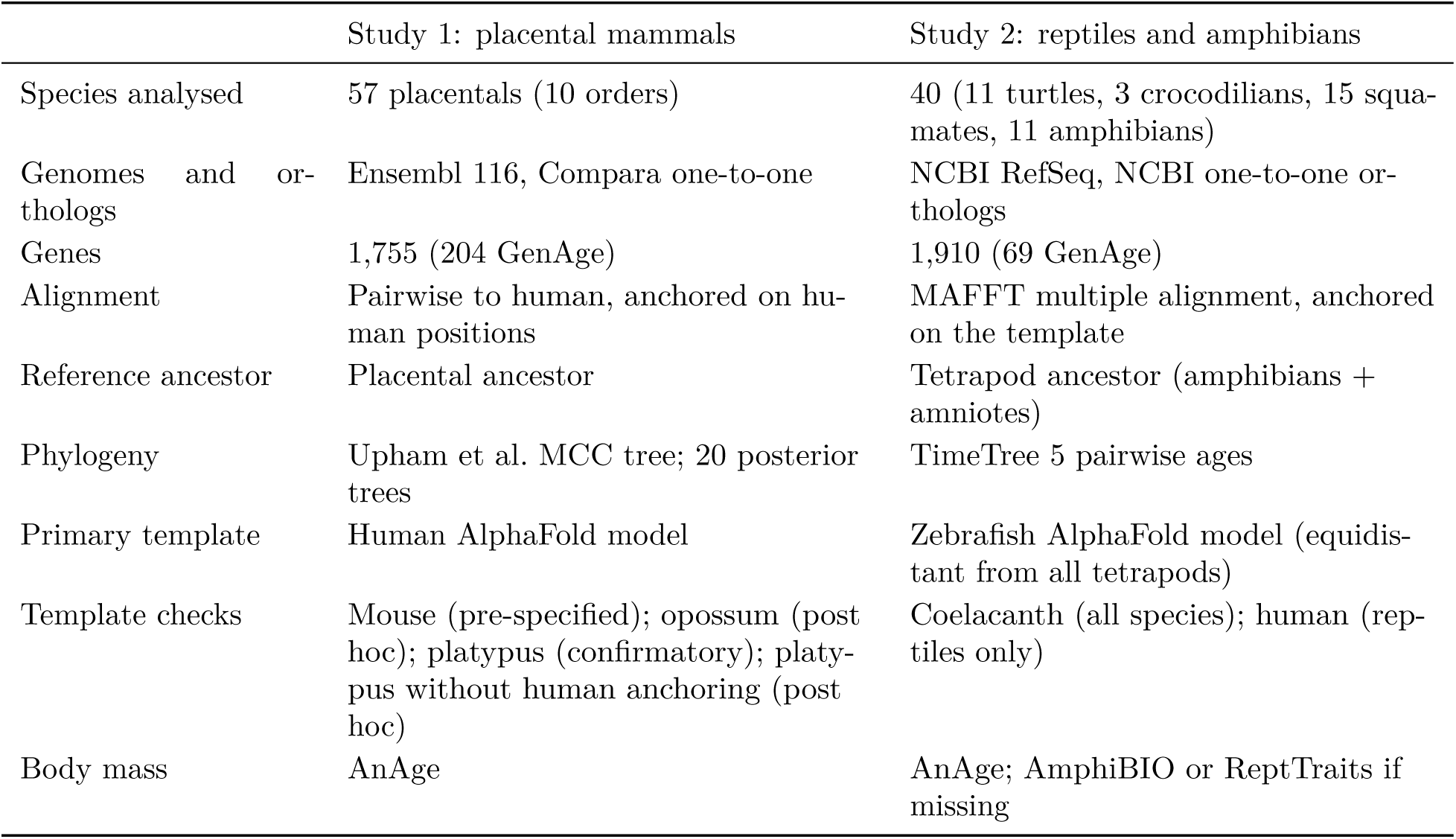
Design of the two studies.

### 2.2 Species and phenotypes

Maximum recorded lifespan and adult mass came from AnAge [12] (quality “acceptable” or “high”, sample size other than “tiny”), matched by binomial or by exact-binomial NCBI Taxonomy synonyms of the same taxon. Relative longevity (LQ) in mammals was lifespan divided by 4.88 *M* ^0.153^ [13]. In Study 1, genomes were the Ensembl 116 mammal assemblies [14] matching AnAge and the dated tree of Upham et al. [15]; eight placentals with early low-coverage assemblies failed the ortholog-coverage filters, leaving 57 species, with six marsupials and monotremes as outgroups. In Study 2, genomes were NCBI-annotated reference assemblies [16] of turtles, crocodilians, lepidosaurs and amphibians; missing masses were taken from AmphiBIO [17] or ReptTraits [18]. Of 48 species with lifespan and mass, 40 passed the coverage filters; the others lacked a RefSeq annotation usable for orthology or, for the allotetraploid *Xenopus laevis*, one-to-one orthologs. Divergence times among these species, zebrafish, spotted gar, coelacanth and human came from the TimeTree 5 API [19]; an average-linkage tree reproduced them exactly (root 435 Ma). Lifespan and body mass of all species analysed are shown in Figure S1 and listed, with their species-level metrics, in Tables S6 and S7.

### 2.3 Genes, alignments and ancestral reconstruction

Each study analysed human genes from GenAge [12] and a background of 2,000 other genes taken in a fixed random order, with proteins of 50–1,000 residues and a full-length AlphaFold DB model [20, 21] identical to the template sequence (human in Study 1, zebrafish in Study 2). In Study 1, orthologs were aligned pairwise to the human protein (BLOSUM62, free end gaps) and anchored on human positions; in Study 2, all orthologs, three fishes and human were aligned with MAFFT [22] and columns were defined by the zebrafish residues. Eligible sites had pLDDT *≥* 70 in the template model; species–gene pairs, genes and species were retained by coverage thresholds fixed in the plans (*≥* 80% of eligible sites per pair, *≥* 60% of species per gene with outgroups present, *≥* 70% of genes per species). Ancestral states were inferred with IQ-TREE 2.4.0 [23] under LG+G4 [24] on the fixed species topology, at the placental (Study 1) or tetrapod (Study 2) ancestor; mean posterior probabilities of the most probable state were 0.992 and 0.937, and the ancestral node was verified independently in 60 random genes per study.

### 2.4 Stability prediction and metrics

ThermoMPNN predicts ΔΔ*G*(*a → b*) at site *i* as *f_i_*(*b*) *− f_i_*(*a*) from frozen ProteinMPNN embeddings of the backbone and full sequence. We threaded the maximum a posteriori ancestral sequence onto the template backbone and obtained *f_i_*for all sites in one pass; our vectorized implementation reproduced the original per-mutation code to within 10*^−^*^6^. Positive values are destabilizing. For species *s* with residue *b* at site *i*,

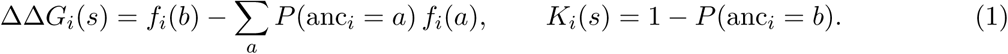

Positions in 10-residue windows with *≥* 5 differences from the ancestor were masked as probable alignment or annotation errors (2.6% of sites in Study 1; 11.8% in Study 2, where many such windows reflect true divergence). The primary metric was *M_s_* = Σ ΔΔ*G/* Σ *K*, the mean predicted ΔΔ*G* per amino-acid difference from the ancestor; secondary metrics were the net ΔΔ*G* per site and the divergence per site, *R_s_* = Σ *K/L*, the expected proportion of sites at which the species differs from the ancestor. A difference is the net outcome of all substitutions at a site along the lineage: a change followed by a reversal leaves no difference, and successive changes count once. *M_s_* and *R_s_* therefore describe net differences rather than every substitution that occurred; with at most about a tenth of the sites differing from the ancestor (mean *R_s_*0.099 in amphibians, the most divergent group), multiple changes at the same site should be uncommon. *M_s_* summarizes how a species’ amino-acid differences from the ancestor are predicted to affect folding when each is evaluated singly, in the ancestral sequence context and on a fixed backbone. It is not a measurement of the stability of the extant proteins: effects are summed additively and epistasis is ignored. For double mutants, additive ThermoMPNN predictions perform comparably to non-additive models, but no current model captures epistasis well [25].

### 2.5 Statistical analysis

In each study, log_10_ maximum lifespan was regressed on the standardized metric and log_10_ adult mass by generalized least squares with Pagel’s *λ* estimated by maximum likelihood [26, 27], with likelihood-ratio tests calibrated by 999 parametric bootstrap simulations re-estimating *λ*. The two primary tests per study (each at *α* = 0.025) were P1 (*M_s_* over all genes) and P2 (*M_s_*over GenAge genes, also compared with 1,000 random background gene sets of equal size). Predictor sanity controls (C1–C3) and a positive control (divergence from the ancestor against lifespan, as a measure of the rate of protein evolution) were specified in advance. Sensitivity analyses are listed in Tables S3 and S4. In the mammal confirmatory amendment E5, all templates were evaluated on identical genes and sites. The primate offset Δ*_T_* was the mean metric of primates minus that of other placentals, in SD, for template *T*. Three hypotheses were fixed with decision rules based on 1,000 bootstrap resamples of genes:

- H5a: Δ_human_ *−* Δ_platypus_ *<* 0;
- H5b: the two outgroup templates agree more with each other than with the human template;
- H5c: P1 with the platypus template gives no association (bootstrap *P ≥* 0.025 and 95% CI including zero).

Per-gene tests were exploratory, with Benjamini–Hochberg correction [28]. After all planned results were known, three post hoc exploratory analyses were specified in dated amendments before being computed (E6 in Study 1, E2 in Study 2; Supplementary Methods). They were a reanalysis of the mammal outgroup template with alignments and sites that did not depend on the human sequence, a follow-up of a nominal human-template association in reptiles, and a calibration of the per-gene tests by simulation under the null hypothesis.

## 3. Results

### 3.1 The predictor captures site-level stability signal in both radiations

Amino-acid differences between the ancestor and extant species had lower predicted ΔΔ*G* than the 19 possible alternatives at the same sites (mammals 0.29 vs 0.64 kcal/mol, 97.3% of genes; reptiles and amphibians 0.28 vs 0.62, 98.3%), as expected under purifying selection. Alternatives were more destabilizing at buried than at exposed sites (1.36 vs 0.60; 1.33 vs 0.57), and proline and glycine were the most destabilizing replacements in helices and strands (Figure 2). These controls show that the predictor is informative at the level of sites; they do not show that differences in *M_s_* between species reflect differences in the stability of extant proteins, nor that the metric is sensitive enough to detect species-level effects of a given size. Species-level values were nevertheless highly reproducible across independent outgroup templates (Sections 3.3 and 3.4).

**Figure 2:**
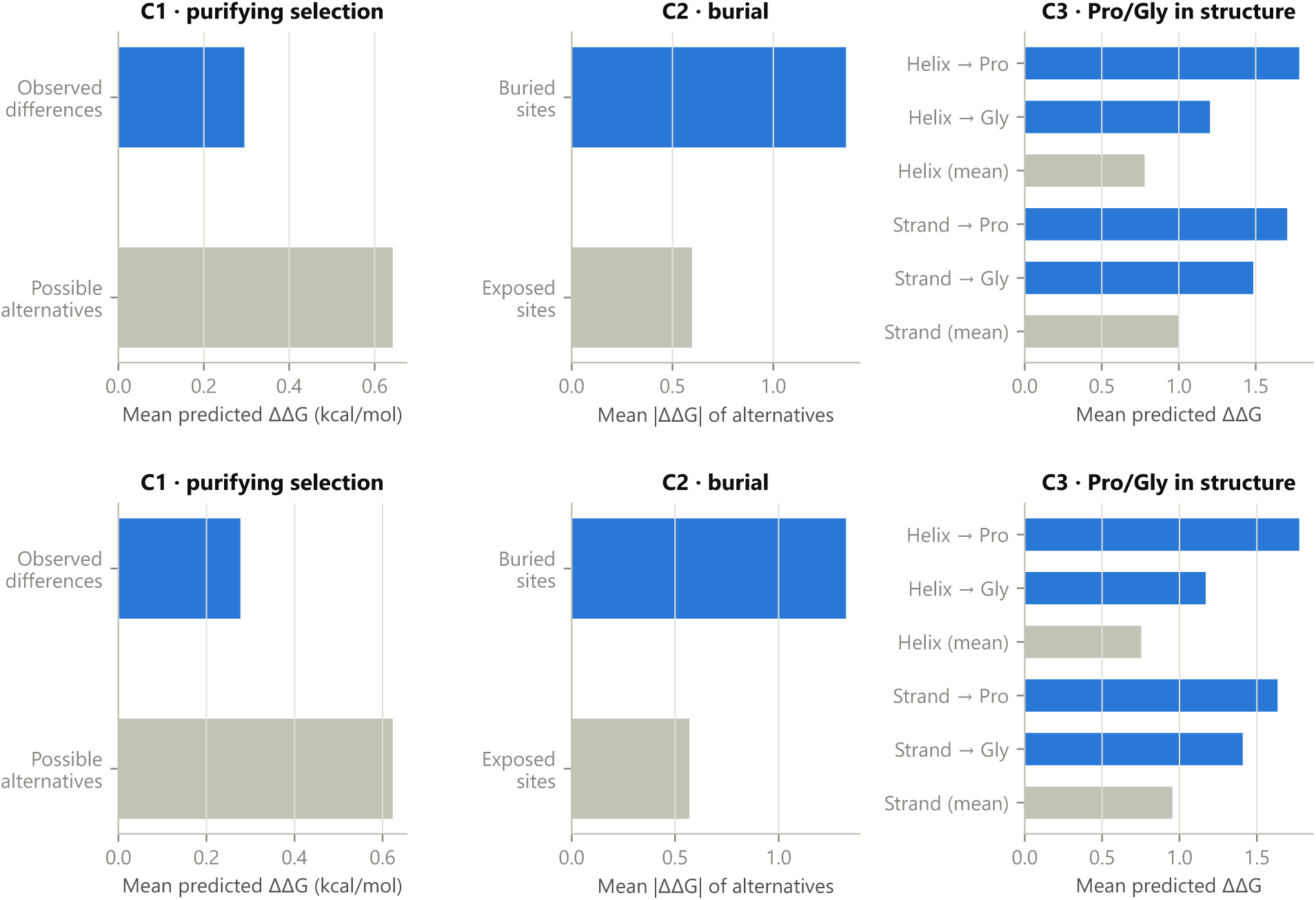
Predictor sanity controls in placental mammals (top; human template, placental ancestor) and in reptiles and amphibians (bottom; zebrafish template, tetrapod ancestor). C1, observed differences from the ancestor vs possible alternatives at the same sites; C2, buried vs exposed sites; C3, proline and glycine replacements in helical and extended regions.

### 3.2 No detectable association between predicted stability and lifespan

No association between mean predicted ΔΔ*G* per difference and lifespan was detected in mammals (*β* = *−*0.075 log_10_ years per SD, 95% CI *−*0.183 to 0.033, bootstrap *P* = 0.178) or in reptiles and amphibians (*β* = *−*0.015, 95% CI *−*0.119 to 0.089, *P* = 0.799; Table 2, Figure 3). Expressed as change in maximum lifespan per SD of the metric, these intervals span *−*34% to +8% in mammals with the human template, *−*9% to +15% with the platypus template on common sites (*−*8% to +17% with the opossum template), and *−*24% to +23% in reptiles and amphibians; they exclude large effects but remain compatible with moderate ones. Human GenAge genes and their orthologs did not differ from random background sets in either study (empirical *P* = 0.774 and 0.775; Figure S2). In reptiles and amphibians, no subset showed an association: reptiles only *P* = 0.16, squamates only *P* = 0.14, amphibians only *P* = 0.76, turtles excluded *P* = 0.37. Of the remaining sensitivity analyses in both studies, only ordinary least squares without phylogeny in mammals reached *P <* 0.05 (*P* = *<*0.001; Tables S3 and S4, Figure S3).

**Figure 3:**
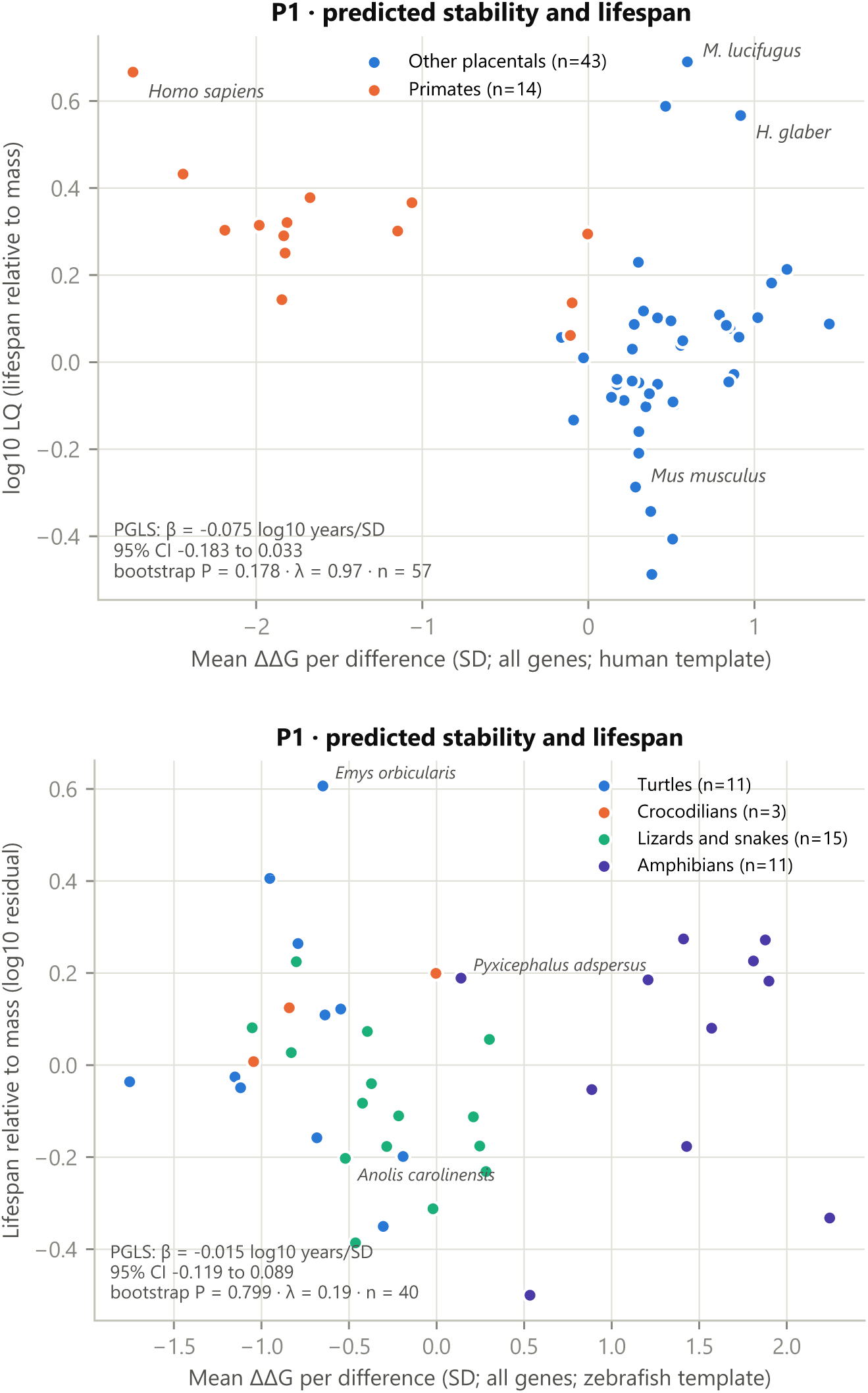
Mean predicted ΔΔ*G* per difference from the ancestor against relative longevity. Top, placental mammals (human template; primates in orange). Bottom, reptiles and amphibians by clade (zebrafish template; lifespan residual on mass shown for display). Insets report the primary PGLS results.

**Table 2:** Primary and secondary tests in both studies. *β* is the change in log_10_ maximum lifespan per SD of the metric, adjusted for log_10_ adult mass. *P* values are parametric-bootstrap likelihood-ratio tests (999 replicates) unless marked.

| Test | $\beta$ | 95% CI | $\lambda$ | $P$ |
| --- | --- | --- | --- | --- |
| *Study 1: placental mammals ($n = 57$)* |  |  |  |  |
| P1: all genes, human template | -0.075 | [-0.183, 0.033] | 0.97 | 0.178 |
| P2: GenAge genes, human template | -0.047 | [-0.126, 0.032] | 0.98 | 0.264 |
| P2 competitive: GenAge vs 1,000 background sets | - | - | - | 0.774<sup>a</sup> |
| Divergence from ancestor per site (secondary) | -0.137 | [-0.252, -0.021] | 0.97 | 0.026 |
| Net $\Delta\Delta G$ per site (secondary) | -0.106 | [-0.216, 0.003] | 0.97 | 0.060 |
| P1, opossum template (post hoc) | 0.014 | [-0.038, 0.067] | 0.97 | 0.595 |
| P2, opossum template (post hoc) | -0.006 | [-0.048, 0.036] | 0.97 | 0.780 |
| P2 competitive, opossum template (post hoc) | - | - | - | 0.750<sup>a</sup> |
| P1, platypus template, common sites (confirmatory, H5c) | 0.010 | [-0.039, 0.060] | 0.97 | 0.701 |
| P2 competitive, platypus template, common sites | - | - | - | 0.906<sup>a</sup> |
| *Study 2: reptiles and amphibians ($n = 40$)* |  |  |  |  |
| P1: all genes | -0.015 | [-0.119, 0.089] | 0.19 | 0.799 |
| P2: GenAge genes | 0.014 | [-0.089, 0.117] | 0.19 | 0.817 |
| P2 competitive: GenAge vs 1,000 background sets | - | - | - | 0.775<sup>a</sup> |
| Divergence from ancestor per site (positive control) | -0.085 | [-0.174, 0.003] | 0.00 | 0.113 |
| Net $\Delta\Delta G$ per site | -0.069 | [-0.172, 0.033] | 0.09 | 0.262 |

<sup>a</sup>Empirical two-sided $P$ against 1,000 random background gene sets of the same size. Opossum-template analyses were added after the primary results were known; platypus-template analyses follow a pre-specified confirmatory amendment and use the sites common to all three templates.

### 3.3 Template proximity drives an apparent association in mammals

With the human backbone, humans had the lowest mean ΔΔ*G* per difference of all 57 mammals, followed by chimpanzee, gorilla and macaques; the primate mean lay 1.97 SD below that of other placentals (Figure 3). Because primates are long-lived for their size, this produced a weak negative slope, which pre-specified sensitivity analyses exposed: excluding primates reversed its sign (*β* = 0.036), excluding humans alone removed it (*β* = *−*0.011, *P* = 0.82), and a mouse backbone reduced it (*−*0.024 to *−*0.004 in 200 genes; Figure S4). After seeing these results we repeated the analysis with an opossum backbone, which is equidistant from all placentals: the slope became *β* = 0.014 (95% CI *−*0.038 to 0.067, *P* = 0.595).

We then specified, before retrieving any of its data, the confirmatory amendment E5 with a platypus backbone (Figure 4). On 1,341 genes and identical sites, the primate offset was *−*1.96 SD with the human template, *−*0.73 with the opossum and *−*0.62 with the platypus. The difference Δ_human_ *−* Δ_platypus_ was *−*1.34 SD (95% CI *−*1.80 to *−*1.04), that is, the primate offset was about two-thirds smaller with the platypus template (H5a confirmed). The two outgroup templates gave almost identical species values (*r* = 0.99), whereas each agreed only moderately with the human template (*r* = 0.56 and 0.62; H5b confirmed), and no association with lifespan was detected (*β* = 0.010, 95% CI *−*0.039 to 0.060, *P* = 0.701; H5c met).

**Figure 4:**
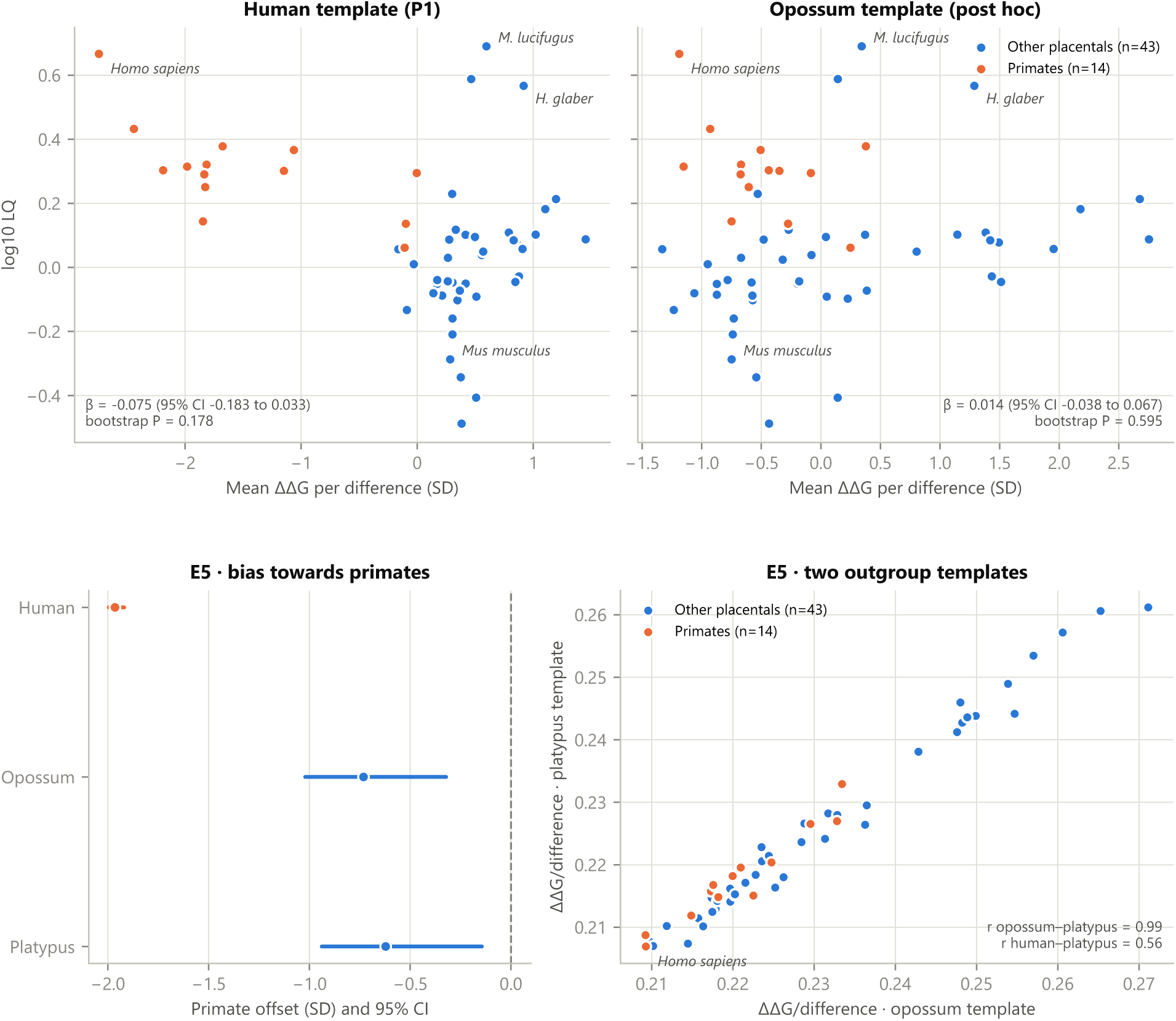
Template bias in mammals. Top, the same species and metric with the human backbone (pre-specified primary analysis) and with the opossum backbone (post hoc). Bottom, confirmatory amendment E5 on identical genes and sites: primate offset Δ*_T_* with each template (95% gene-bootstrap intervals; H5a) and agreement between the opossum and platypus templates (H5b).

A smaller primate offset and the lowest value for humans persisted with both outgroup templates. Both outgroup analyses shared features that could favour humans: eligible sites were defined on the human model and alignments were anchored on human positions. In a post hoc analysis (amendment E6) we removed both. We realigned the orthologs of 1,321 of the 1,341 E5 genes with MAFFT, kept the columns of the platypus protein, defined eligible sites on the platypus model alone and reconstructed the placental ancestor again, so that no step used the human sequence or model as a reference. The primate offset was essentially unchanged: *−*0.52 SD (95% CI *−*0.87 to *−*0.09), against *−*0.60 SD for the human-anchored platypus analysis on the same genes (difference 0.07, 95% CI *−*0.08 to 0.22). Species values were nearly identical (*r* = 0.993). Humans were no longer the lowest: counting from the lowest value they ranked 4 of 57 (2 with the human-anchored analysis on these genes), and no association with lifespan was detected (*β* = 0.013, 95% CI *−*0.035 to 0.062, *P* = 0.56). Restricting the new analysis to sites that were also eligible in the human model changed the offset by only *−*0.09 SD. Human anchoring therefore does not explain the residual primate offset. Whether it reflects a genuine property of primate proteins or another bias remains open.

### 3.4 Equidistant in time may not be equidistant in sequence: templates in reptiles

In Study 2 the zebrafish template was chosen to be equidistant from all species. A second fish, the coelacanth, gave almost identical species values in 300 random genes (*r* = 0.97) and the same null result (*β* = 0.031 and 0.029). A human backbone is also equidistant in time from all reptiles, but within reptiles it separated slowly evolving turtles and crocodilians from faster-evolving squamates more sharply than the zebrafish backbone (Figure 5) and gave a nominal negative slope (*β* = *−*0.130, analytic Wald *P* = 0.012, with *λ* estimated at 0.00), against *−*0.047 (*P* = 0.42) with zebrafish on the same genes. A *λ* near zero does not by itself make an association spurious, so we examined this slope in post hoc analyses (amendment E2; Table S5). A parametric bootstrap likelihood-ratio test, which accounts for the uncertainty in *λ*, gave *P* = 0.088. Among reptiles the human-template metric was strongly correlated with divergence from the ancestor (*r* = 0.89, against 0.68 with zebrafish), and so was the difference between the two templates (*r* = 0.50). Adjusting for divergence halved the slope (*−*0.060, *P* = 0.53). With a clade covariate (turtles and crocodilians against squamates) the slope was similar (*−*0.116), but its standard error tripled (0.150; *P* = 0.45), because the metric itself largely separates the clades; these data cannot distinguish a lifespan effect from a clade effect. Within squamates the slope was *−*0.037 (*P* = 0.35, *n* = 15). Within turtles and crocodilians it was *−*0.120 (*P* = 0.036, *n* = 14; Wald test, not bootstrapped). By the criterion fixed in the amendment the result is therefore mixed: the slope lost nominal significance with the clade covariate and within squamates, but did not approach zero with the covariate and remained nominal within turtles and crocodilians. With the zebrafish template no analysis approached significance. We do not regard the human-template slope as evidence of a lifespan association. Its dependence on divergence fits a mechanism in which lineages that have changed less since the ancestor remain more compatible with a given backbone, so that a template equidistant in time can still favour slowly evolving clades.

**Figure 5:**
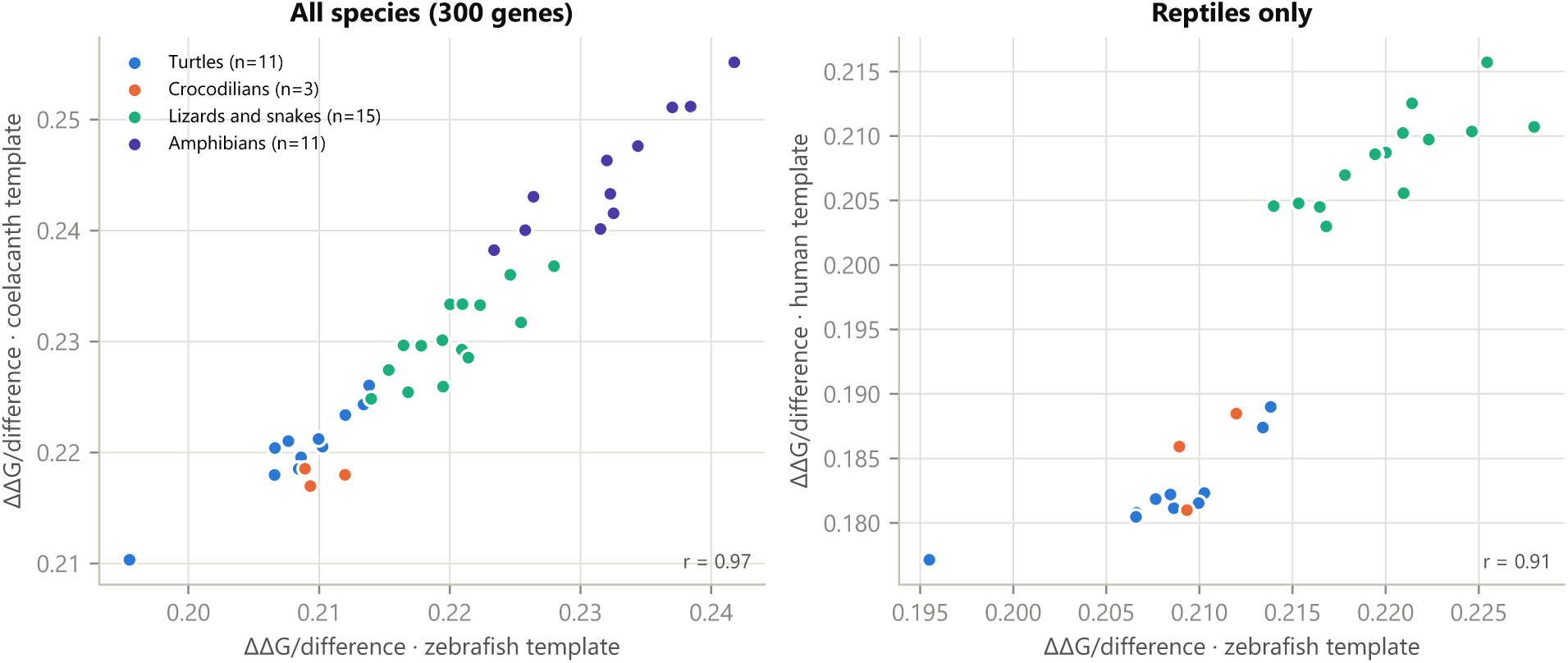
Templates in reptiles and amphibians (300 random genes). Left, coelacanth vs zebrafish backbones for all species. Right, human vs zebrafish backbones for reptiles only.

### 3.5 Clade patterns and secondary analyses

In mammals, divergence per site from the ancestor, the positive control, was lower in longer-lived species (*β* = *−*0.137, 95% CI *−*0.252 to *−*0.021, bootstrap *P* = 0.026), reproducing the known association between longevity and slower molecular evolution [29]. In reptiles and amphibians the direction was the same but not significant (*β* = *−*0.085, *P* = 0.113); turtles and crocodilians had the lowest divergence (0.083 and 0.083 expected differences per site, vs 0.094 in squamates and 0.099 in amphibians), consistent with their slowly evolving genomes [30, 31]. The largest clade difference in the stability metric was between amphibians and reptiles (+1.88 SD for amphibians), which does not follow lifespan, since amphibian lifespans lie between those of squamates and turtles; turtles differed only modestly from squamates (*−*0.51 SD). The metric was not associated with body mass in either study (mammals *P* = 0.63; reptiles and amphibians *P* = 0.12), so the nearly neutral prediction was not supported either.

Per-gene tests were exploratory. In mammals, one of 1,478 genes passed a false discovery rate of 5% (NAT14, a background gene, in the direction opposite to the stability hypothesis); in reptiles and amphibians, none of 1,798 did (Figure S5). In Study 2, 207 genes had nominal *P <* 0.05 (90 expected), with 50.6% of slopes positive. A post hoc calibration (amendment E2) accounted for this excess. We simulated lifespans 1,000 times under the null model and repeated the same per-gene tests. A mean of 139 genes then had *P <* 0.05 (95% interval 25–294; false-positive rate 7.7%), so the observed number is compatible with no association (calibrated *P* = 0.23; Figure S6). The average rate was inflated because many per-gene fits estimated *λ* near zero (38% below 0.05), and the interval was wide because all genes are tested against the same lifespans. With *λ* fixed at the null-model value the tests were calibrated (null false-positive rate 5.0%), and 111 genes were nominal (null mean 90, 95% interval 25–214). Against the simulated distributions, 131 genes had empirical *P <* 0.05, a number within that null range, and none passed the false discovery threshold.

## 4. Discussion

In two independent radiations, endothermic placental mammals and ectothermic reptiles and amphibians, we did not detect an association between longevity and the mean predicted effect of the amino-acid differences that species have accumulated relative to a reconstructed ancestor, and human GenAge genes and their orthologs behaved like random background genes. In both studies the predictor distinguished observed from possible amino-acid changes and buried from exposed sites, so the predictor is not uninformative. These controls do not, however, establish that it can resolve species-level differences in the stability of extant proteins, so the null results speak to the predicted effects of these differences rather than to measured stability.

The confidence intervals limit, but do not exclude, an association. Per SD of the metric, they span *−*9% to +15% of maximum lifespan in mammals with an outgroup template and *−*24% to +23% in reptiles and amphibians, so moderate effects remain compatible with the data. For reference, the minimum effects detectable with 80% power at the two-sided *α* = 0.025 of the primary tests, computed as 3.08 standard errors of *β*, were about 20% (mammals, opossum template) and 44% (reptiles and amphibians). Thousands of genes improve the precision of each species’ estimate, but the comparative sample sizes remain 57 and 40 species. Effects confined to particular proteins or regulatory regions, and properties other than folding stability (activity, aggregation, turnover, modification), also remain open. Turtles, the clearest example of slow ageing among ectotherms [2], did not show a distinctive folding-stability signature; their most distinctive molecular trait in our data was slow evolution.

Folding stability of isolated chains is only one layer of proteostasis. Many protein complexes are conserved across animals [32], but orthology does not guarantee that every interaction is conserved, and conservation of interactions decreases with sequence divergence [33]; subunits are also often stabilized by assembly with their partners [34]. Our metric scores each protein as a monomer, so residues at interaction interfaces, which are usually solvent-exposed in monomer models, contribute little, and compensatory substitutions between interacting partners are invisible. Longevity-related adaptation could therefore act on interfaces, complex assembly or regulation without changing the mean folding effect of amino-acid differences. A natural extension is to separate differences at known or predicted interfaces from those in the structural core, using complex structures rather than monomers and without assuming that the human interactome applies unchanged to all species.

The methodological findings may be the most broadly useful. Structure-based predictors evaluate amino-acid changes in the context of one backbone, and that backbone carries the imprint of the sequences it was predicted for. In mammals, a human backbone made primates, which are long-lived for their size, appear least destabilized, which mimicked support for the stability hypothesis. In reptiles, a human backbone, although equidistant in time, appeared to favour slowly evolving and long-lived turtles and crocodilians, and the metric it produced tracked divergence from the ancestor. Farré et al. [5] compared FoldX energies of modelled human and rat proteins for 40 pairs and larger reference sets, with a different metric and design. Our results do not show that their conclusion is biased, but they indicate that comparisons of this kind should be checked for sensitivity to the structural template. We suggest four checks for comparative structure-based analyses:

1. measure amino-acid differences from a reconstructed ancestor, not from a modern reference species;
2. use a template outside the clade analysed and confirm results with a second, unrelated outgroup template;
3. test whether apparent associations follow the clade of the template species or slowly evolving lineages;
4. test how much an association depends on contrasts between a few clades, for example with bootstrap tests, clade covariates or within-clade analyses, particularly when *λ* is estimated near zero.

### Limitations

ThermoMPNN predictions for single mutations are noisy and were summed additively, ignoring epistasis, protein–protein interactions, oligomers, ligands and modifications [25]; the metric was not validated against measured stability differences between species. Ancestors were reconstructed, with greater uncertainty at the tetrapod node. Study 1 used human-anchored pairwise alignments and human-defined eligible sites; a post hoc reanalysis without them did not change the residual primate offset, whose origin remains unexplained. Study 2 used multiple alignments but masked more sites, partly true divergence. Species were sampled by genome availability (57 and 40), few lineages are exceptionally long-lived, AnAge records only maximum lifespans, masses in Study 2 mix typical and maximum values, and environmental temperature, which may affect both stability and ageing in ectotherms, could not be adjusted because it was available for only 69% of reptiles, below the pre-specified threshold. The follow-up of the human template in reptiles, the calibration of the per-gene tests and the human-independent mammal reanalysis were specified after the planned results were known and are exploratory; with 29 reptiles in two main clades, a clade effect and a lifespan effect cannot be separated. The opossum analysis was post hoc; its main conclusion was confirmed under a confirmatory amendment, but neither plan was deposited in a public registry, and the plan hashes document content rather than timing.

## 5. Conclusion

In two independent tetrapod radiations, the choice of structural template was enough to create an apparent association between predicted protein stability and longevity. A template from within the sample favoured the clade of the template species, and a template equidistant in time still favoured slowly evolving lineages. Ancestral references, outgroup templates and explicit checks for clade and rate effects should therefore be standard in comparative structure-based studies. We demonstrated the bias for ThermoMPNN applied to single-chain models; whether other predictors are equally sensitive remains to be tested. With these safeguards, in the proteins and 57 + 40 species analysed, we detect no robust association between longevity and the mean predicted effect on folding stability of amino-acid differences from a reconstructed ancestor; moderate effects cannot be excluded.

## Declarations

### Data and code availability

All code, both analysis plans with their dated amendments and SHA256 hashes, the provenance log of every download (URL, retrieval date and SHA256), the result tables and the generated inputs of this manuscript are available at https://github.com/Enso-bio/tetrapod-template-bias. Downloaded inputs and large intermediate files (alignments, ancestral reconstructions and ThermoMPNN energies) are regenerated by the scripts, and TimeTree divergence times are re-queried rather than redistributed. The plan hashes verify integrity; independent evidence of when the plans were written would require a public registry.

## Supplementary material

### Supplementary Methods

The tables and values in this section were generated from the project files (recorded package versions, download logs and code) rather than typed by hand. Every download was logged with its URL, UTC retrieval time and the SHA256 of the stored file. Both studies ran in the same Python environment (Windows-11-10.0.26200). Software and versions are listed in Table S1, and data sources with their retrieval dates in Table S2.

**Table S1:** Software and versions.

| Software | Version | Use |
| --- | --- | --- |
| Python | 3.12.10 | All analyses |
| NumPy [35] | 2.5.3 | Numerical computation, PGLS |
| SciPy [36] | 1.18.1 | Optimisation of $\lambda$, distributions |
| pandas [37] | 3.0.6 | Data handling |
| statsmodels [38] | 0.15.0 | OLS with HC3 errors (sensitivity) |
| Biopython [39] | 1.88 | Pairwise alignment, structures, trees, accessibility |
| PyTorch [40] | 2.14.1 (CUDA 13.0) | ThermoMPNN inference (NVIDIA GeForce RTX 5070) |
| OmegaConf | 2.3.1 | ThermoMPNN configuration |
| openpyxl | 3.1.5 | Reading ReptTraits (Study 2) |
| matplotlib [41] | 3.11.2 | Figures |
| ThermoMPNN [10] | Kuhlman-Lab main branch, downloaded 8 Oct 2026 | Stability prediction (weights thermoMPNN_default.pt) |
| ProteinMPNN [42] | Weights v_48_020, bundled with ThermoMPNN | Frozen backbone and sequence embeddings |
| IQ-TREE [23] | 2.4.0 | Ancestral sequence reconstruction |
| MAFFT [22] | 7.526 (Windows package) | Multiple alignments (Study 2) |

#### Genes and orthologs

In both studies the gene universe was restricted to human proteins of 50–1,000 residues. GenAge human genes were always included, and background genes were taken in a fixed random order (NumPy generator, seed 20261008 in Study 1 and 20261009 in Study 2) until 2,000 passed the template criteria. Study 1 used the `ortholog_one2one` homologies of the Ensembl 116 Compara files and the protein listed for each homology. The human template was the AlphaFold DB model whose sequence was identical to that protein. Study 2 used the `Ortholog` pairs relative to human in the NCBI `gene_orthologs` file. For each gene it took the MANE Select or RefSeq Select protein when annotated, and the longest RefSeq protein otherwise. Genes required exactly one zebrafish ortholog; the zebrafish template was the longest isoform identical in sequence to a UniProtKB entry with a full-length AlphaFold DB model. Genome assemblies and annotations are listed in Tables S8 and S9.

#### Alignment and coverage

In Study 1, each ortholog was aligned to the human protein with the Biopython `PairwiseAligner` in global mode with free end gaps. Scoring used BLOSUM62, gap opening *−*10 and extension *−*0.5, and the ortholog residue at each human position was retained. In Study 2, the orthologs, three fishes and human were aligned with MAFFT (`--auto`, one thread), and the columns with a zebrafish residue were retained. The same pairwise settings were used to map residues onto the alternative templates (mouse, opossum, platypus, coelacanth and human). A species–gene pair was valid if *≥* 80% of the eligible template sites (pLDDT *≥* 70) were aligned. A gene was retained if it was valid in *≥* 60% of the analysed species and also present in at least one of the six marsupial and monotreme outgroups (Study 1) or in *≥* 2 of the 3 fishes (Study 2). Species were retained if they were valid in *≥* 70% of the retained genes.

**Table S2:** Data sources. Retrieval dates are UTC ranges over both studies; the Study column gives the studies that used each source.

| Source | Version or release | Retrieved | Study |
| --- | --- | --- | --- |
| AnAge (HAGR) | Build 15 (released 3 Jul 2023) | 8–9 Oct 2026 | 1, 2 |
| GenAge, human genes (HAGR) | Build 21 (released 28 Aug 2023) | 8–9 Oct 2026 | 1, 2 |
| Ensembl | Release 116; REST and FTP (Compara homologies, peptides, genome list) | 8 Oct 2026 | 1 |
| UniProtKB | Release 2026_03 | 8–9 Oct 2026 | 1, 2 |
| AlphaFold DB | Model version 6 (all templates used) | 8–9 Oct 2026 | 1, 2 |
| MamPhy (Upham et al.) | GitHub commit 740b701; node-dated MCC tree and 100 posterior trees | 8 Oct 2026 | 1 |
| NCBI Taxonomy (E-utilities) | Current at retrieval | 8–9 Oct 2026 | 1, 2 |
| NCBI Datasets | RefSeq genome annotations (Table S9) | 9 Oct 2026 | 2 |
| NCBI gene_orthologs | FTP file, Last-Modified 9 Oct 2026 | 9 Oct 2026 | 2 |
| TimeTree 5 | Pairwise API (adjusted divergence times) | 9 Oct 2026 | 2 |
| AmphiBIO | v1 (figshare file 8828578) | 9 Oct 2026 | 2 |
| ReptTraits | v1-2 (figshare file 45408133) | 9 Oct 2026 | 2 |

#### Trees and ancestral reconstruction

In Study 1 the node-dated MCC tree of Upham et al. [15] was pruned to the analysed species and outgroups, and Nexus annotations were removed. In Study 2 the tree was the average-linkage (UPGMA) tree of TimeTree pairwise ages; its maximum deviation from the queried ages was below 10*^−^*^12^ Ma. For each gene the tree was pruned to the species present, and IQ-TREE was run as `-te` *tree* `-m LG+G4 -asr -nt 1 -seed` *seed*. The topology was fixed and the branch lengths were re-estimated. The ancestral node (placental or tetrapod ancestor) was the most recent common ancestor of the ingroup in the IQ-TREE output tree rooted on the outgroups. In 60 random genes per study this node was identified independently by re-rooting with Biopython, and all agreed. The maximum a posteriori ancestral sequence was threaded onto the template, and the full posterior distribution of ancestral states entered ΔΔ*G_i_*(*s*) and *K_i_*(*s*).

#### Structural quantities and ThermoMPNN

pLDDT values were read from the B-factor column of each AlphaFold model. Solvent accessibility was computed on the template model with the Biopython Shrake–Rupley algorithm [43] (100 points per atom, probe radius 1.4 Å). It was normalised by the theoretical maximum accessibilities of Tien et al. [44], and sites with relative accessibility *<* 0.25 were classified as buried. ThermoMPNN used the published default weights (file `thermoMPNN_default.pt`, SHA256 `af449118ccfb4e34971d802321907ed82a8a035ae80f055b f8fc56009a4a838a`) on frozen ProteinMPNN v_48_020 embeddings. The model configuration was the published one: the embeddings of two ProteinMPNN layers were concatenated and passed through light attention and an output network with hidden layers of 64 and 32 units. The threaded sequence replaced the model sequence while the backbone coordinates were kept; selenocysteine was scored as cysteine. A single forward pass returned the 20-value energy vector *f_i_* for every site. On test proteins, the resulting *f_i_*(*b*) *− f_i_*(*a*) matched the original per-mutation code to within 10*^−^*^6^ kcal/mol. The window mask excluded all positions of any 10-residue window containing *≥* 5 differences between the species and the ancestor.

#### Statistics

The phylogenetic covariance *C* was computed from the most recent common ancestor of the analysed species and scaled to a mean root-to-tip distance of 1. Pagel’s *λ* transformation was *V* (*λ*) = *λC* + (1 *− λ*) diag(*C*). Generalized least squares used the eigen-decomposition of the correlation matrix, so that each value of *λ* required only a weighted least-squares fit. The profile likelihood of *λ* was evaluated on a 41-point grid over [0, 1]. It was then refined by bounded Brent optimisation (tolerance 10*^−^*^4^) between the neighbours of the best grid point, keeping the better of the two values. Confidence intervals and Wald tests used the *t* distribution with *n − p* degrees of freedom. For the bootstrap likelihood-ratio tests, 999 data sets were simulated under the fitted null model (coefficients, residual variance and *λ*). Both models were refitted to each data set with *λ* re-estimated, and *P* = (1 + #*{*LR*^∗^ ≥* LR*}*)*/*(*B* + 1). The competitive test recomputed the metric for 1,000 random sets of background genes of the same size as the GenAge set and refitted the PGLS model, re-estimating *λ*, for each set. The two-sided empirical *P* was (1 + #*{|β^∗^| ≥ |β*_GenAge_*|}*)*/*1001. The predictor was standardized to unit SD before fitting. Posterior-tree sensitivity in Study 1 used 20 trees evenly spaced among the 100 downloaded posterior trees. Each analysis used a seed derived from the study seed, and rerunning the metric and statistics steps reproduced all results exactly.

#### Post hoc analyses

These analyses were written into the analysis plans as dated amendments (E6 in Study 1, E2 in Study 2), and the plans were hashed before any of them was computed. In E6 the input genes were the 1,341 common genes of E5. For each gene the Compara one-to-one proteins of the 57 placentals and six outgroups, with human as one more species, were aligned with MAFFT (--auto). The columns with a platypus residue were kept, and eligible sites were those with pLDDT *≥* 70 in the platypus model. The coverage rules were those of the main analysis, with the gene filter applied to the 57 placentals and no refiltering of species; 1,321 genes were retained. The placental ancestor was reconstructed with the same IQ-TREE settings (mean posterior of the most probable state 0.974), threaded onto the platypus model and scored with ThermoMPNN and the same window mask. Primate offsets were compared with those of E5 on the same genes by 1,000 paired bootstrap resamples of genes (seed 20261070), and P1 was tested with 999 bootstrap simulations (seed 20261071). In E2(a), the species metrics of the template sensitivity analysis (300 genes, 29 reptiles) were used. They were analysed with bootstrap likelihood-ratio tests (999 simulations; seeds 20261080 and 20261081), with a clade covariate (turtles and crocodilians against squamates), within clades, and with adjustment for log_10_ divergence from the ancestor over all genes (Table S5). In E2(b), the null model (log_10_ lifespan on log_10_ mass; *λ* = 0.18) was fitted to the 40 species. 1,000 responses were simulated from it (seed 20261082), and every per-gene test was repeated on each with the same species, *λ* estimated by maximum likelihood and the Wald test. A variant fixed *λ* at the null-model value. Empirical per-gene *P* values were (1 + #*{P ^∗^ ≤ P }*)*/*(*B* + 1).

**Figure S1:**
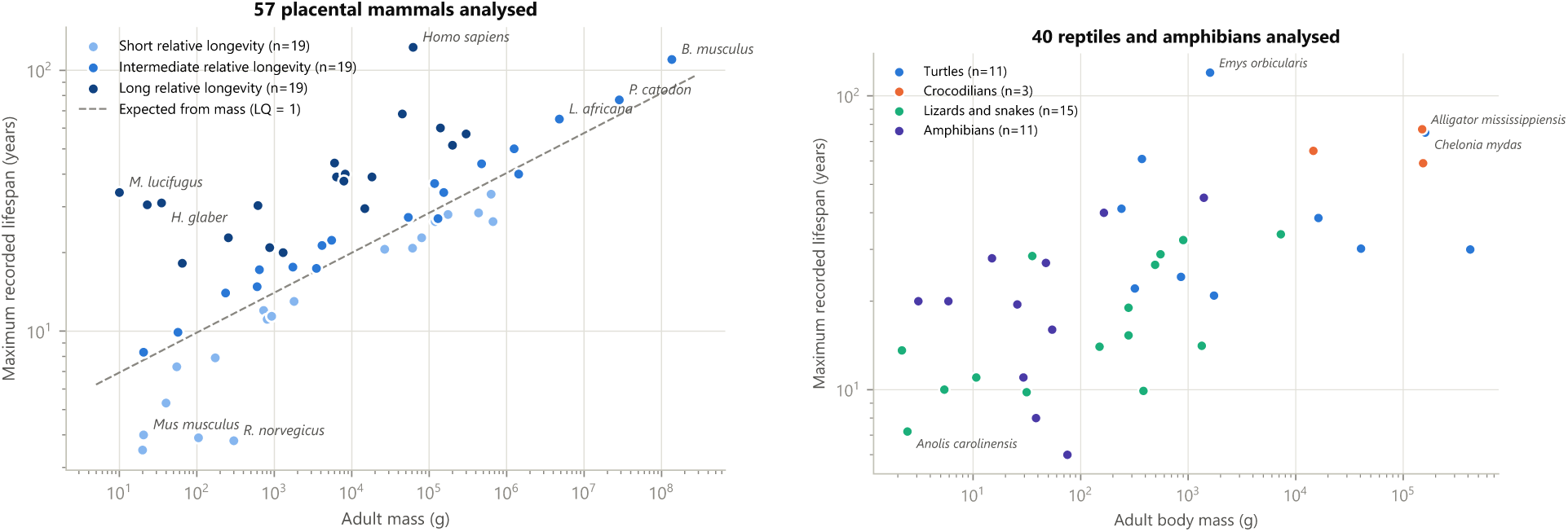
Species analysed: maximum recorded lifespan against adult mass. Left, placental mammals by LQ tertile. Right, reptiles and amphibians by clade.

**Figure S2:**
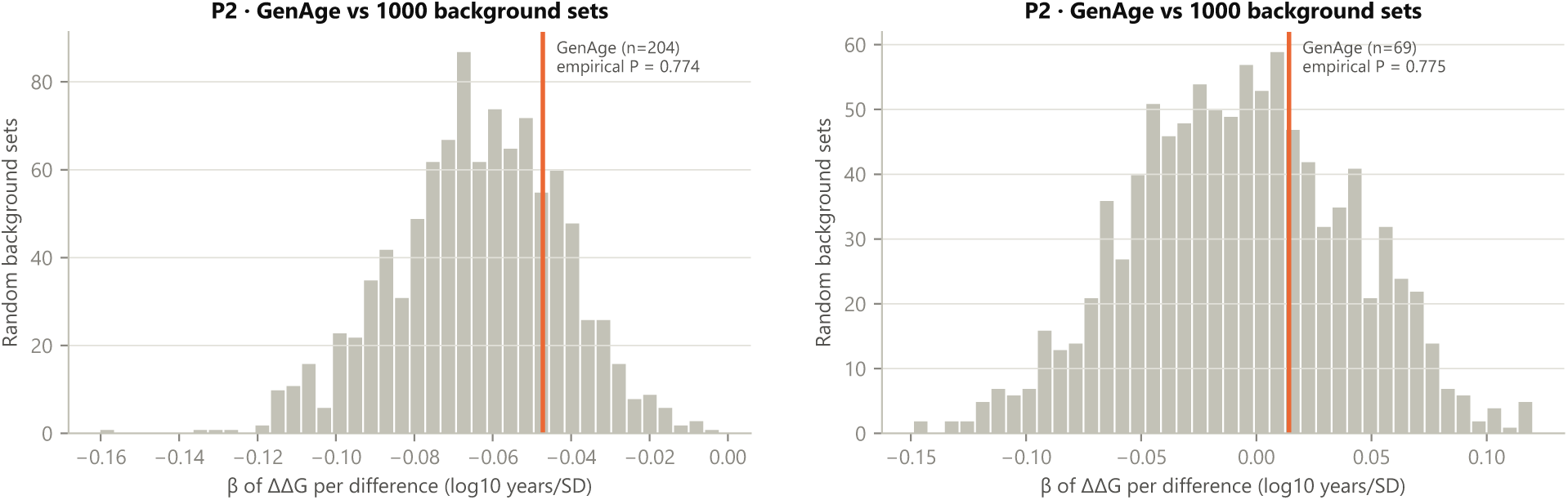
Competitive tests of human GenAge genes and their orthologs against 1,000 random background gene sets of equal size. Left, mammals; right, reptiles and amphibians.

**Figure S3:**
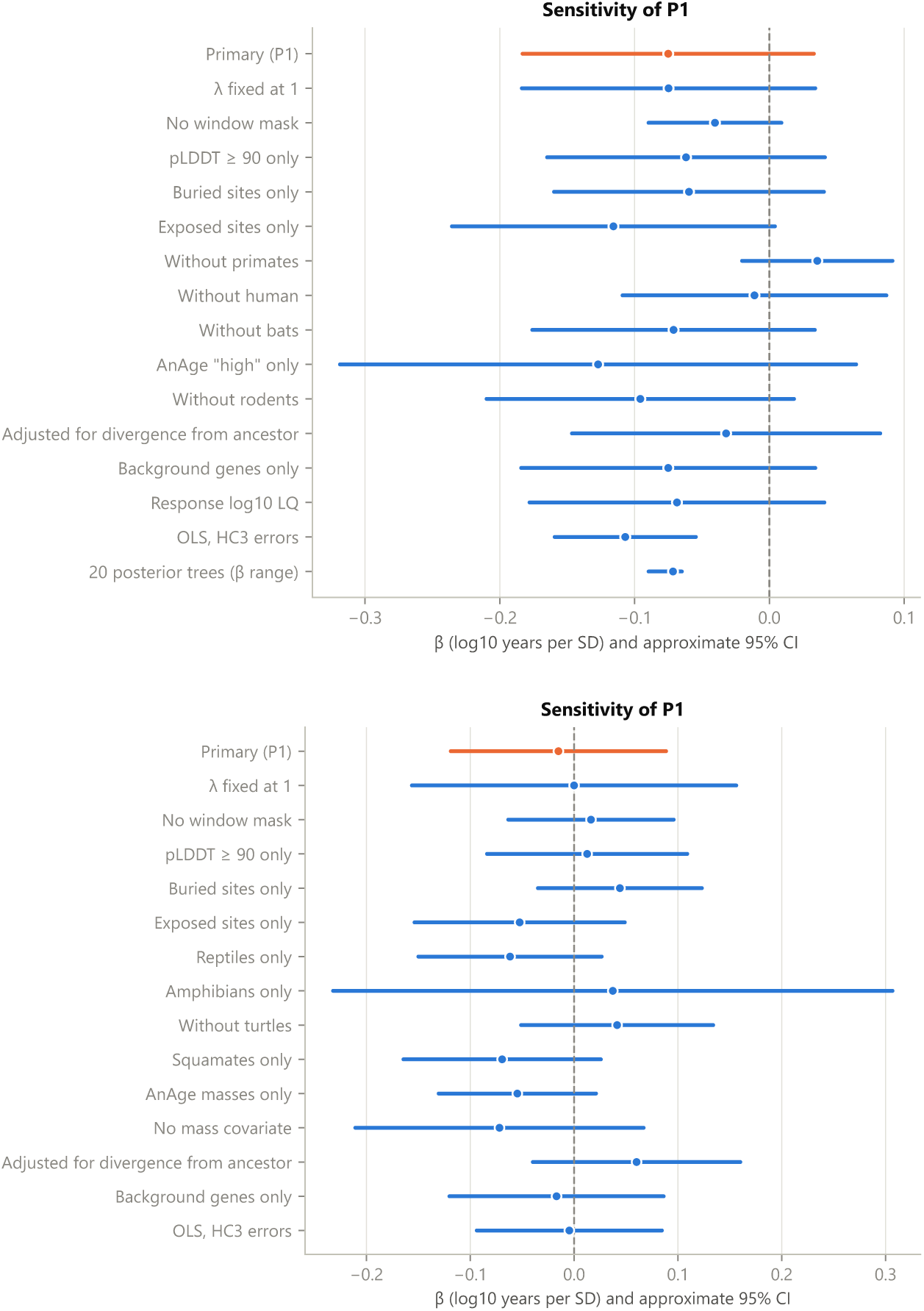
Sensitivity analyses of P1. Top, mammals (last row: range over 20 posterior trees); bottom, reptiles and amphibians.

**Figure S4:**
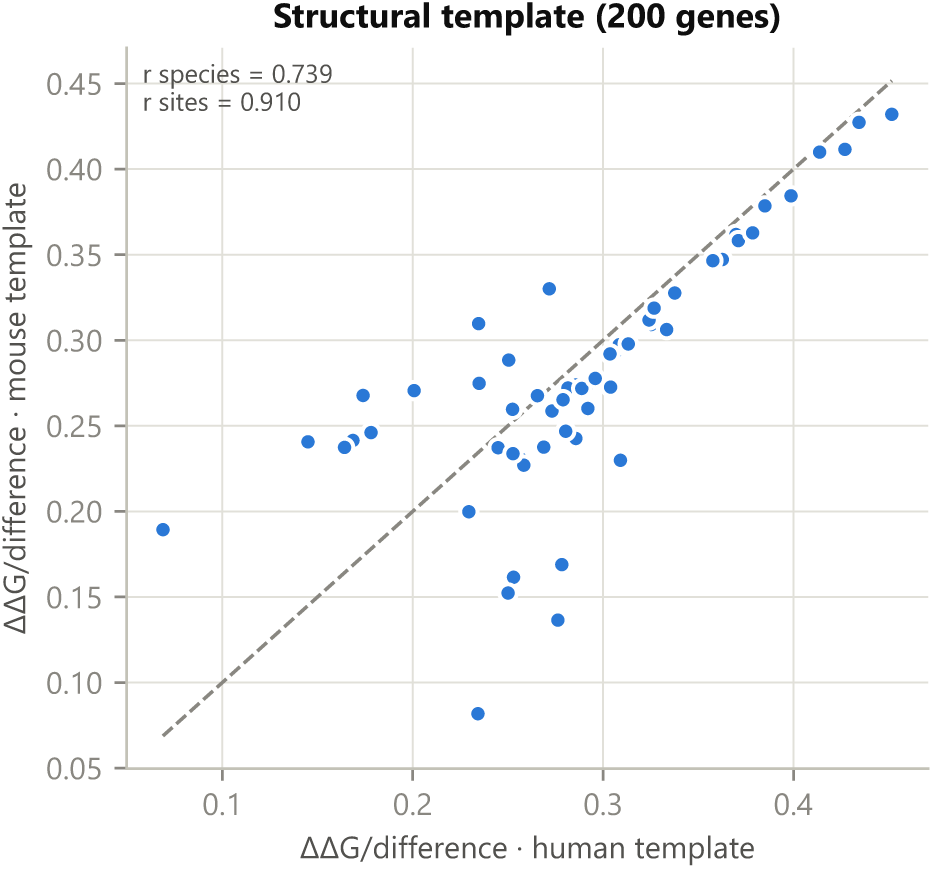
Mammals: species-level metric with human and mouse backbones in 200 random genes (pre-specified template sensitivity).

**Figure S5:**
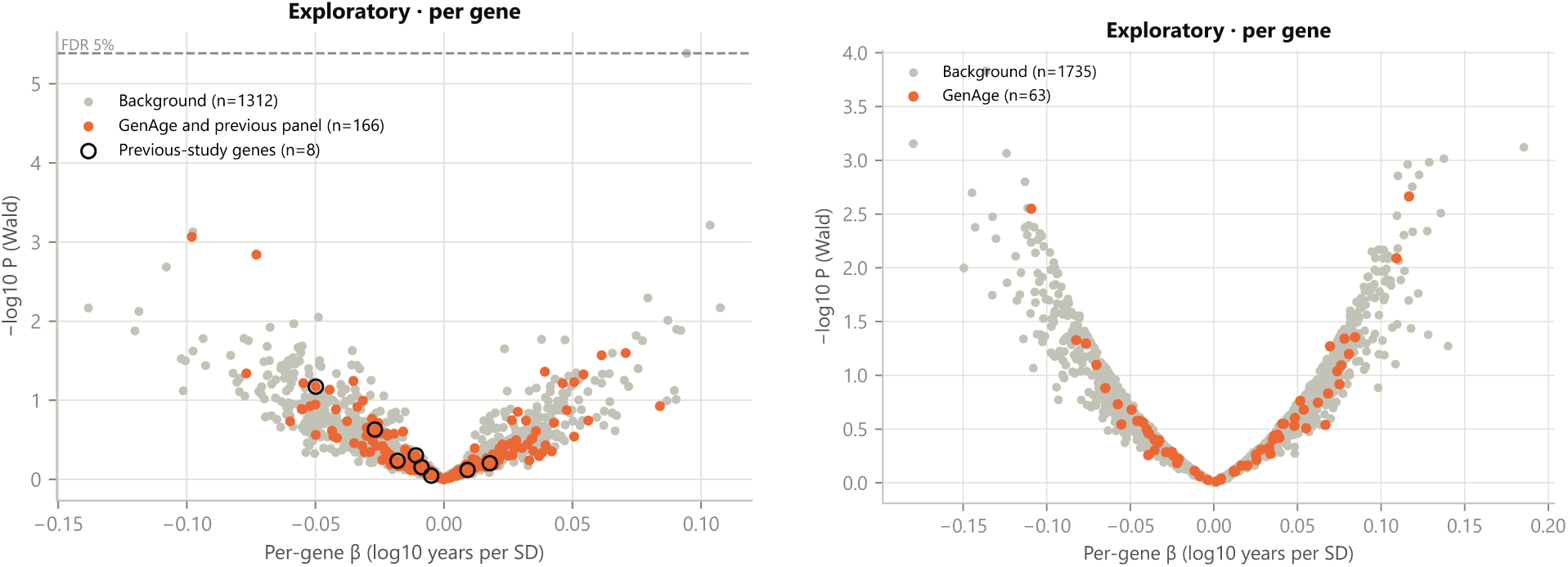
Exploratory per-gene tests. Left, mammals (rings: genes from a previous candidate panel); right, reptiles and amphibians.

**Figure S6:**
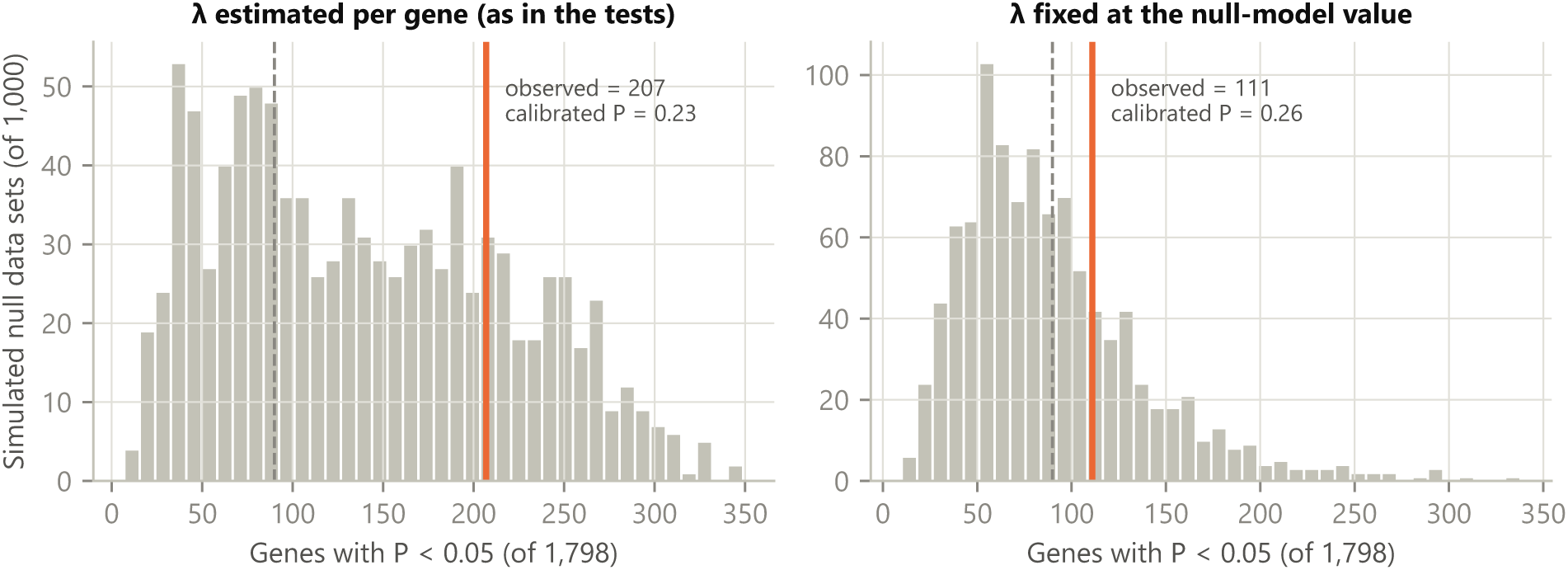
Post hoc calibration of the per-gene tests in reptiles and amphibians (amendment E2). Distribution of the number of genes with *P <* 0.05 in 1,000 data sets simulated under the null model, with *λ* estimated per gene (left) or fixed at the null-model value (right). Orange line, observed number; dashed line, 5% of the tests.

**Table S3:** Sensitivity analyses of P1 in mammals (analytic Wald tests conditional on *λ*; diagnostics, not discovery tests).

| Analysis | $n$ | $\beta$ | 95% CI | $\lambda$ | $P$ |
| --- | --- | --- | --- | --- | --- |
| $\lambda$ fixed at 1 | 57 | -0.075 | [-0.184, 0.034] | 1.00 | 0.17 |
| No window mask | 57 | -0.040 | [-0.090, 0.009] | 0.99 | 0.11 |
| Sites with pLDDT $\geq 90$ | 57 | -0.062 | [-0.165, 0.041] | 0.97 | 0.23 |
| Buried sites only | 57 | -0.060 | [-0.160, 0.040] | 0.97 | 0.24 |
| Exposed sites only | 57 | -0.116 | [-0.236, 0.004] | 0.98 | 0.058 |
| Primates excluded | 43 | 0.036 | [-0.020, 0.091] | 0.98 | 0.21 |
| *Homo sapiens* excluded | 56 | -0.011 | [-0.109, 0.087] | 0.99 | 0.82 |
| Bats excluded | 54 | -0.071 | [-0.176, 0.034] | 0.94 | 0.18 |
| AnAge quality “high” only | 15 | -0.127 | [-0.319, 0.064] | 0.97 | 0.17 |
| Rodents excluded | 42 | -0.096 | [-0.210, 0.018] | 0.93 | 0.097 |
| Adjusted for $\log_{10}$ divergence from ancestor | 57 | -0.032 | [-0.146, 0.082] | 0.97 | 0.58 |
| Background genes only | 57 | -0.075 | [-0.184, 0.034] | 0.97 | 0.17 |
| Response $\log_{10}$ LQ (no mass term) | 57 | -0.069 | [-0.178, 0.041] | 0.97 | 0.21 |
| OLS with HC3 errors (no phylogeny) | 57 | -0.107 | [-0.159, -0.055] | – | <0.001 |
| 20 posterior trees (range) | 57 | -0.090 to -0.065 | – | – | 0.12–0.25 |

**Table S4:**
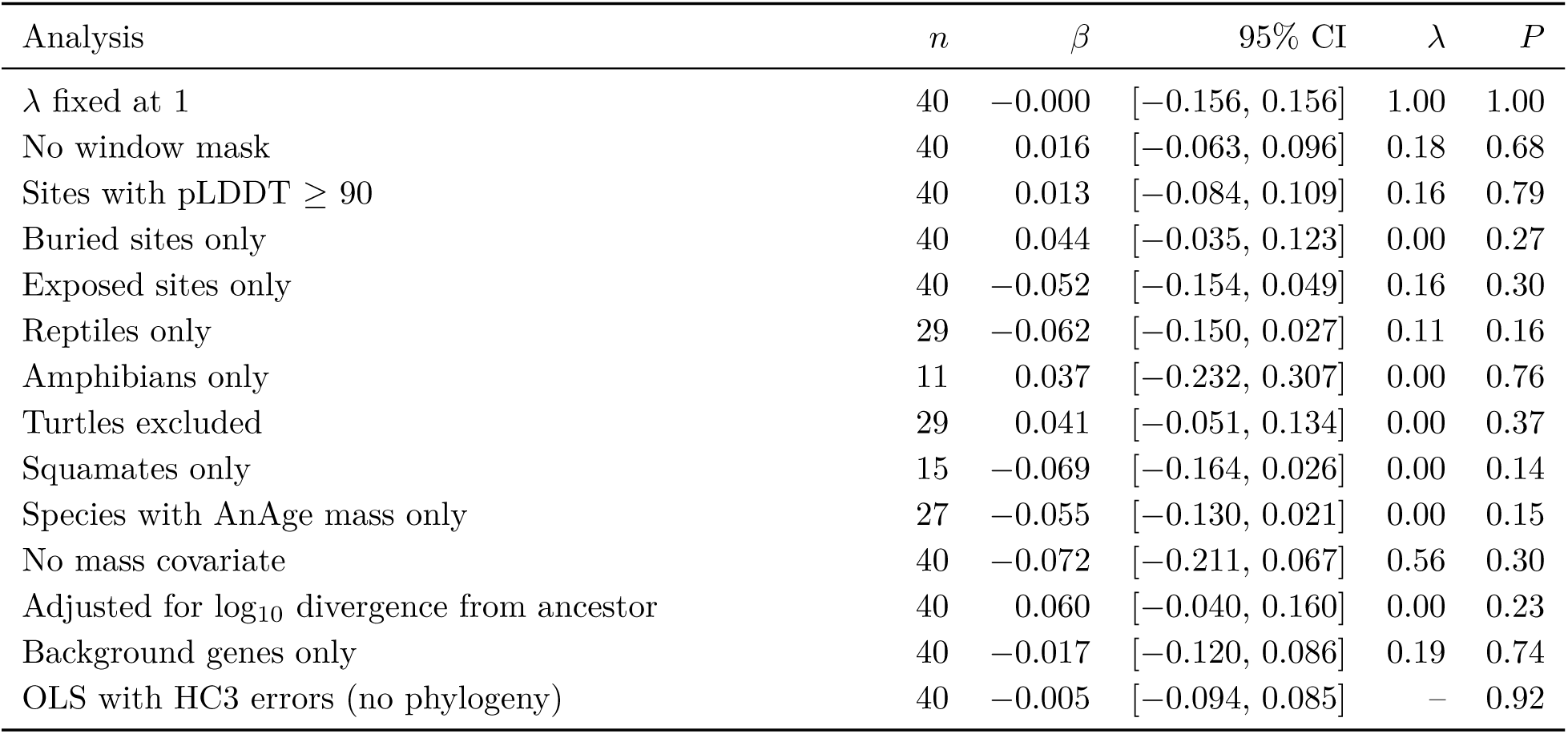
Sensitivity analyses of P1 in reptiles and amphibians (analytic Wald tests conditional on *λ*).

**Table S5:**
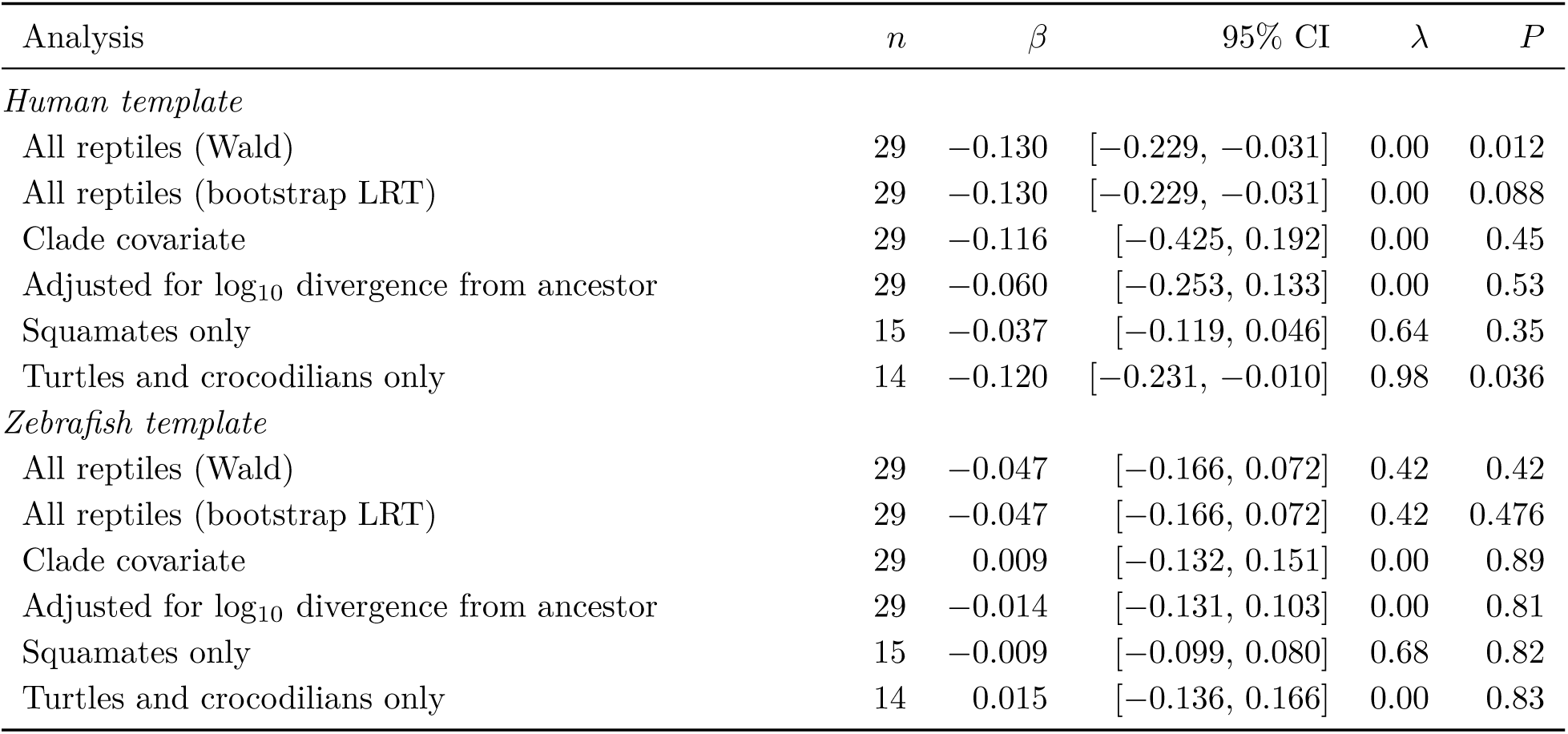
Post hoc follow-up of the human-template association in reptiles (amendment E2; 300 genes, sites shared by the zebrafish, coelacanth and human templates). Wald tests conditional on *λ* unless stated.

**Table S6:** Mammal species (names as in AnAge). Lifespan in years; mass in g; group, LQ tertile (S, I, L); *M*_hum_ and *M*_opo_, mean predicted ΔΔ*G* per difference from the ancestor with human and opossum templates; *R*, expected proportion of sites differing from the placental ancestor.

| Species | Order | Lifespan | Mass | LQ | Group | $M_{\text{hum}}$ | $M_{\text{opo}}$ | $R$ |
| --- | --- | --- | --- | --- | --- | --- | --- | --- |
| *Bison bison* | Artiodactyla | 33.5 | 630,000 | 0.89 | S | 0.229 | 0.223 | 0.0441 |
| *Bos grunniens* | Artiodactyla | 26.3 | 667,000 | 0.69 | S | 0.224 | 0.215 | 0.0454 |
| *Camelus dromedarius* | Artiodactyla | 28.4 | 434,000 | 0.80 | S | 0.235 | 0.229 | 0.0408 |
| *Capra hircus* | Artiodactyla | 20.8 | 61,000 | 0.79 | S | 0.226 | 0.217 | 0.0456 |
| *Ovis aries* | Artiodactyla | 22.8 | 80,000 | 0.83 | S | 0.215 | 0.210 | 0.0444 |
| *Sus scrofa* | Artiodactyla | 27 | 130,000 | 0.91 | I | 0.217 | 0.214 | 0.0413 |
| *Ailuropoda melanoleuca* | Carnivora | 36.8 | 117,500 | 1.26 | I | 0.229 | 0.231 | 0.0385 |
| *Canis lupus* | Carnivora | 20.6 | 26,625 | 0.89 | S | 0.217 | 0.213 | 0.0377 |
| *Mustela putorius* | Carnivora | 11.1 | 809 | 0.82 | S | 0.219 | 0.217 | 0.0403 |
| *Neovison vison* | Carnivora | 11.4 | 925 | 0.82 | S | 0.215 | 0.213 | 0.0396 |
| *Panthera leo* | Carnivora | 28 | 175,000 | 0.90 | S | 0.222 | 0.223 | 0.0368 |
| *Panthera pardus* | Carnivora | 27.3 | 53,750 | 1.06 | I | 0.221 | 0.221 | 0.0368 |
| *Panthera tigris* | Carnivora | 26.3 | 119,700 | 0.90 | S | 0.251 | 0.248 | 0.0379 |
| *Ursus americanus* | Carnivora | 34 | 154,250 | 1.12 | I | 0.237 | 0.238 | 0.0380 |
| *Ursus maritimus* | Carnivora | 43.8 | 475,000 | 1.21 | I | 0.250 | 0.247 | 0.0390 |
| *Vulpes vulpes* | Carnivora | 21.3 | 4,132 | 1.22 | I | 0.222 | 0.218 | 0.0380 |
| *Balaenoptera musculus* | Cetacea | 110 | 136,000,000 | 1.28 | I | 0.248 | 0.246 | 0.0410 |
| *Delphinapterus leucas* | Cetacea | 40 | 1,430,000 | 0.94 | I | 0.252 | 0.247 | 0.0415 |
| *Monodon monoceros* | Cetacea | 50 | 1,250,000 | 1.20 | I | 0.251 | 0.248 | 0.0414 |
| *Physeter macrocephalus* | Cetacea | 77 | 28,500,000 | 1.14 | I | 0.254 | 0.255 | 0.0412 |
| *Tursiops truncatus* | Cetacea | 51.6 | 200,000 | 1.63 | L | 0.269 | 0.266 | 0.0427 |
| *Myotis lucifugus* | Chiroptera | 34 | 10 | 4.90 | L | 0.238 | 0.231 | 0.0473 |
| *Pteropus vampyrus* | Chiroptera | 20.9 | 872 | 1.52 | L | 0.264 | 0.258 | 0.0439 |
| *Rhinolophus ferrumequinum* | Chiroptera | 30.5 | 23 | 3.87 | L | 0.232 | 0.228 | 0.0422 |
| *Dasypus novemcinctus* | Cingulata | 22.3 | 5,500 | 1.22 | I | 0.281 | 0.267 | 0.0430 |
| *Oryctolagus cuniculus* | Lagomorpha | 13 | 1,800 | 0.85 | S | 0.227 | 0.231 | 0.0457 |
| *Equus caballus* | Perissodactyla | 57 | 300,000 | 1.70 | L | 0.224 | 0.218 | 0.0370 |
| *Callithrix jacchus* | Primates | 22.8 | 255 | 2.00 | L | 0.151 | 0.220 | 0.0422 |
| *Gorilla gorilla* | Primates | 60.1 | 139,842 | 2.01 | L | 0.098 | 0.219 | 0.0392 |
| *Homo sapiens* | Primates | 122.5 | 62,035 | 4.64 | L | 0.070 | 0.208 | 0.0381 |
| *Macaca fascicularis* | Primates | 39 | 6,362 | 2.09 | L | 0.117 | 0.216 | 0.0387 |
| *Macaca mulatta* | Primates | 40 | 8,235 | 2.06 | L | 0.109 | 0.208 | 0.0379 |
| *Macaca nemestrina* | Primates | 37.6 | 7,912 | 1.95 | L | 0.116 | 0.216 | 0.0389 |
| *Mandrillus leucophaeus* | Primates | 39 | 18,250 | 1.78 | L | 0.116 | 0.217 | 0.0393 |
| *Microcebus murinus* | Primates | 18.2 | 65 | 1.97 | L | 0.208 | 0.224 | 0.0395 |
| *Nomascus leucogenys* | Primates | 44.1 | 6,000 | 2.39 | L | 0.124 | 0.231 | 0.0399 |
| *Otolemur garnettii* | Primates | 20 | 1,300 | 1.37 | L | 0.203 | 0.221 | 0.0428 |
| *Pan troglodytes* | Primates | 68 | 44,984 | 2.71 | L | 0.086 | 0.212 | 0.0384 |
| *Prolemur simus* | Primates | 17.6 | 1,738 | 1.15 | I | 0.203 | 0.229 | 0.0386 |
| *Rhinopithecus roxellana* | Primates | 29.5 | 14,750 | 1.39 | L | 0.115 | 0.214 | 0.0393 |
| *Saimiri boliviensis* | Primates | 30.3 | 615 | 2.32 | L | 0.155 | 0.218 | 0.0427 |
| *Loxodonta africana* | Proboscidea | 65 | 4,800,000 | 1.27 | I | 0.260 | 0.243 | 0.0386 |
| *Cavia porcellus* | Rodentia | 12 | 728 | 0.90 | S | 0.224 | 0.217 | 0.0543 |
| *Chinchilla lanigera* | Rodentia | 17.2 | 642 | 1.31 | I | 0.225 | 0.222 | 0.0519 |
| *Dipodomys ordii* | Rodentia | 9.9 | 57 | 1.09 | I | 0.236 | 0.224 | 0.0528 |
| *Heterocephalus glaber* | Rodentia | 31 | 35 | 3.69 | L | 0.255 | 0.245 | 0.0553 |
| *Ictidomys tridecemlineatus* | Rodentia | 7.9 | 173 | 0.74 | S | 0.204 | 0.207 | 0.0426 |
| *Jaculus jaculus* | Rodentia | 7.3 | 55 | 0.81 | S | 0.234 | 0.226 | 0.0551 |
| *Marmota marmota* | Rodentia | 17.4 | 3,500 | 1.02 | I | 0.207 | 0.211 | 0.0433 |
| *Mesocricetus auratus* | Rodentia | 3.9 | 105 | 0.39 | S | 0.234 | 0.228 | 0.0571 |
| *Microtus ochrogaster* | Rodentia | 5.3 | 40 | 0.62 | S | 0.224 | 0.215 | 0.0576 |
| *Mus musculus* | Rodentia | 4 | 20 | 0.52 | S | 0.223 | 0.214 | 0.0575 |
| *Mus spicilegus* | Rodentia | 3.5 | 20 | 0.45 | S | 0.227 | 0.217 | 0.0582 |
| *Octodon degus* | Rodentia | 14 | 235 | 1.24 | I | 0.233 | 0.226 | 0.0564 |
| *Peromyscus maniculatus* | Rodentia | 8.3 | 20 | 1.07 | I | 0.222 | 0.216 | 0.0560 |
| *Rattus norvegicus* | Rodentia | 3.8 | 300 | 0.33 | S | 0.228 | 0.219 | 0.0588 |
| *Sciurus vulgaris* | Rodentia | 14.8 | 600 | 1.14 | I | 0.200 | 0.206 | 0.0424 |

**Table S7:**
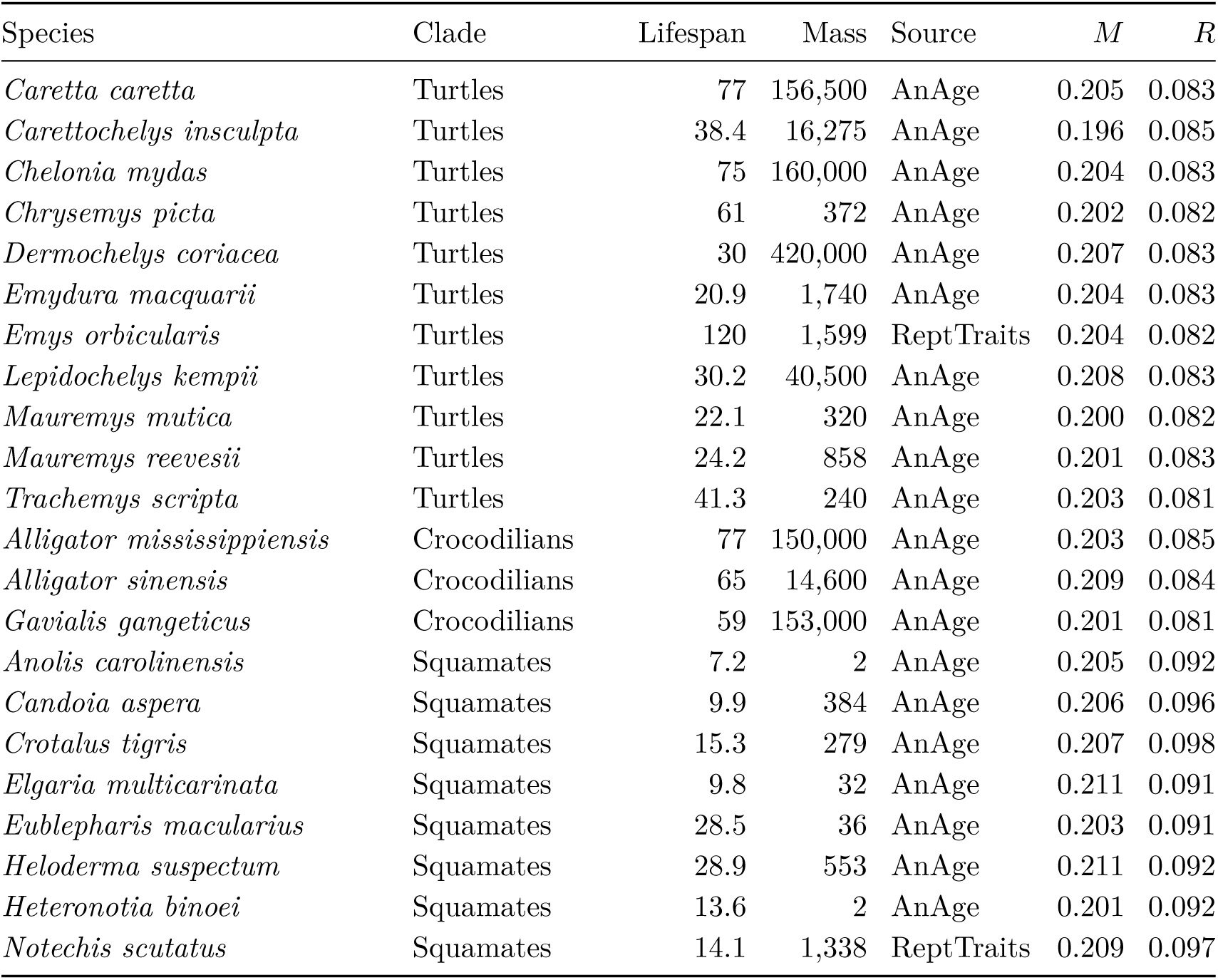
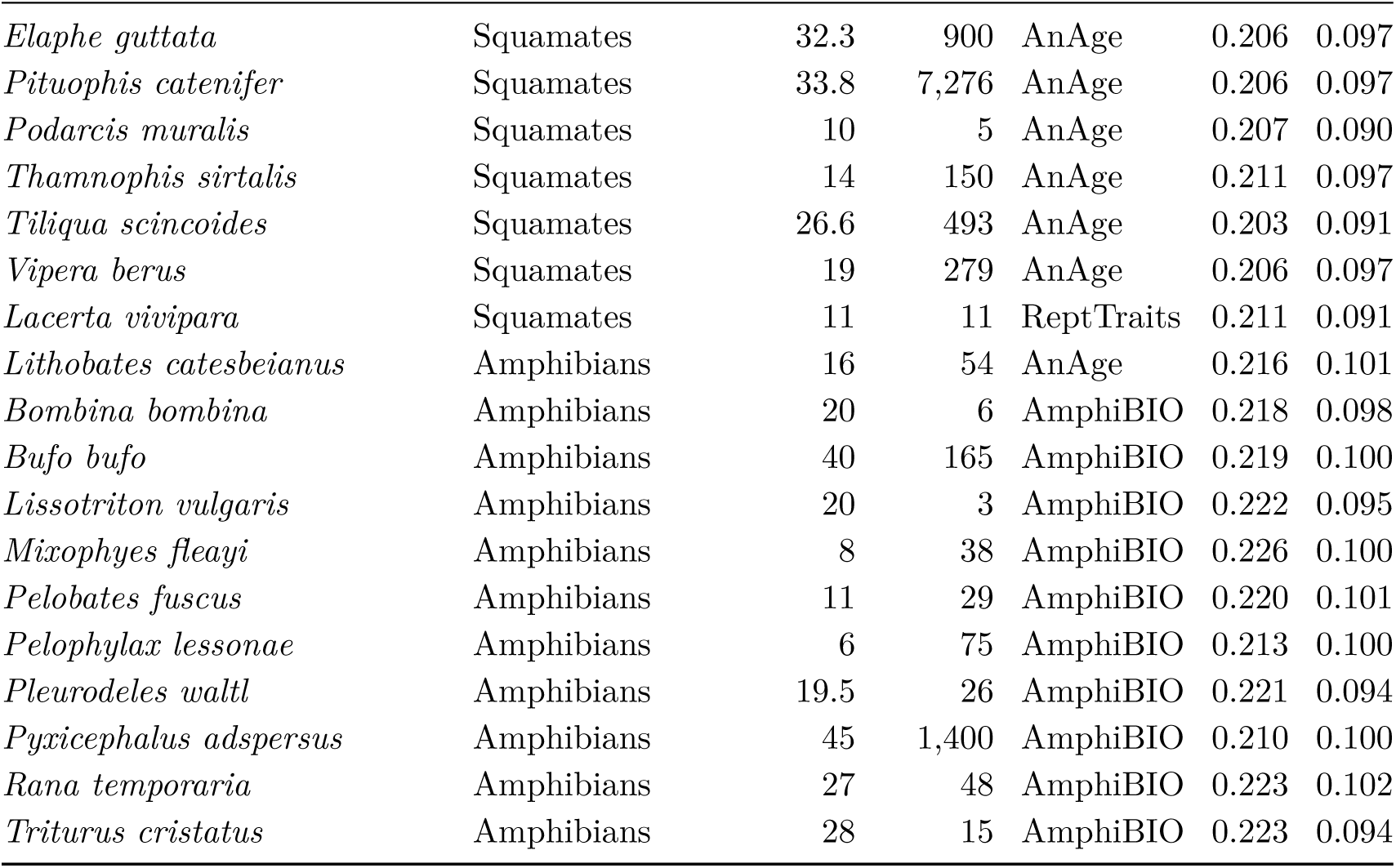
Reptile and amphibian species (names as in AnAge). Lifespan in years; mass in g and its source; *M*, mean predicted ΔΔ*G* per difference from the ancestor (zebrafish template); *R*, expected proportion of sites differing from the tetrapod ancestor.

**Table S8:**
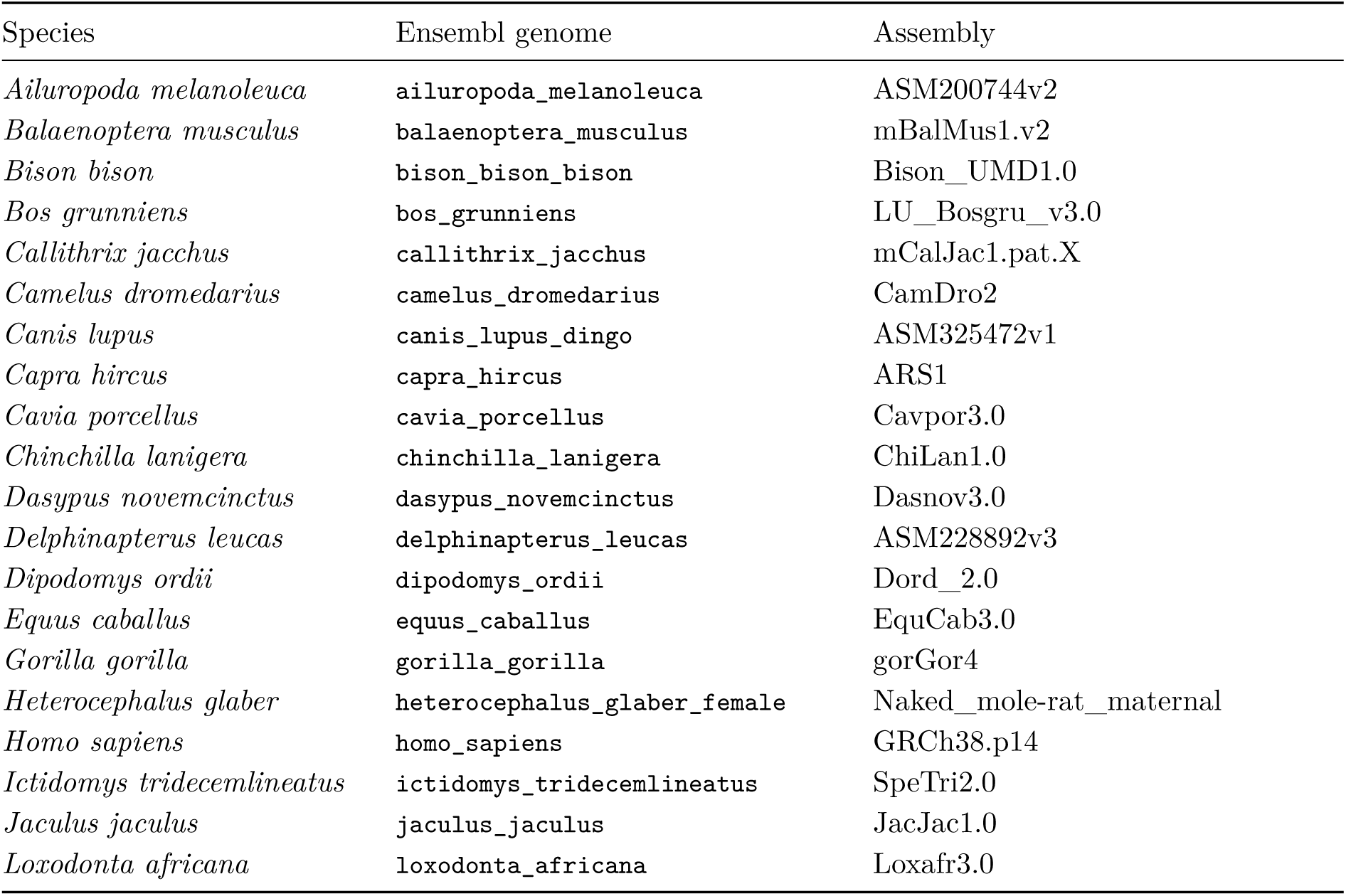
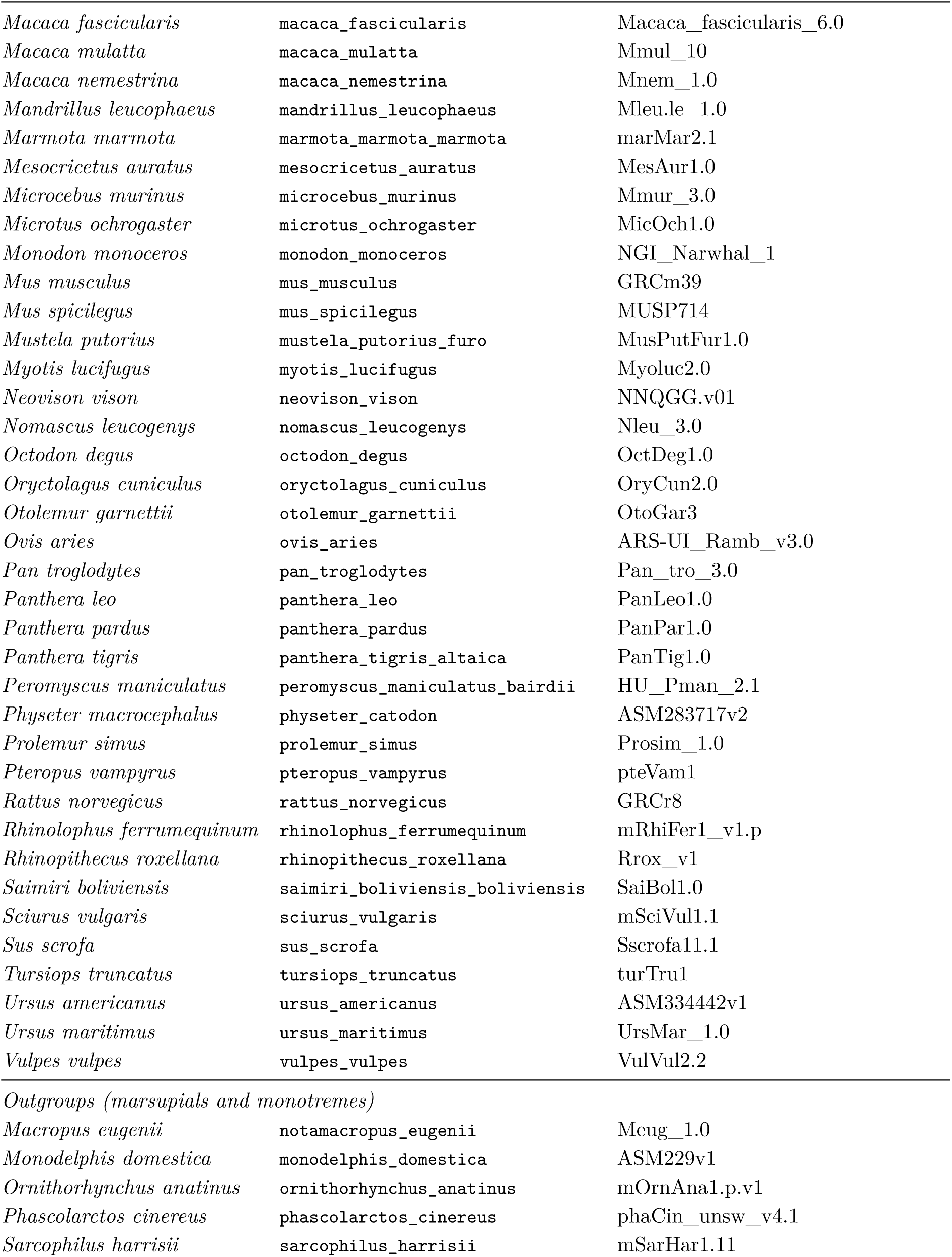
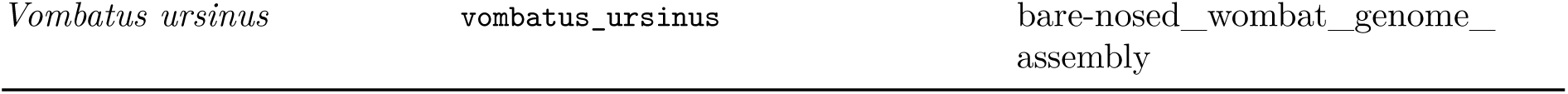
Study 1 genome assemblies (Ensembl 116). Species are named as in AnAge and Table S6; the Ensembl genome is the one used, which for some species is a subspecies or a specific individual. Outgroups were used to root the trees and reconstruct the placental ancestor.

**Table S9:**
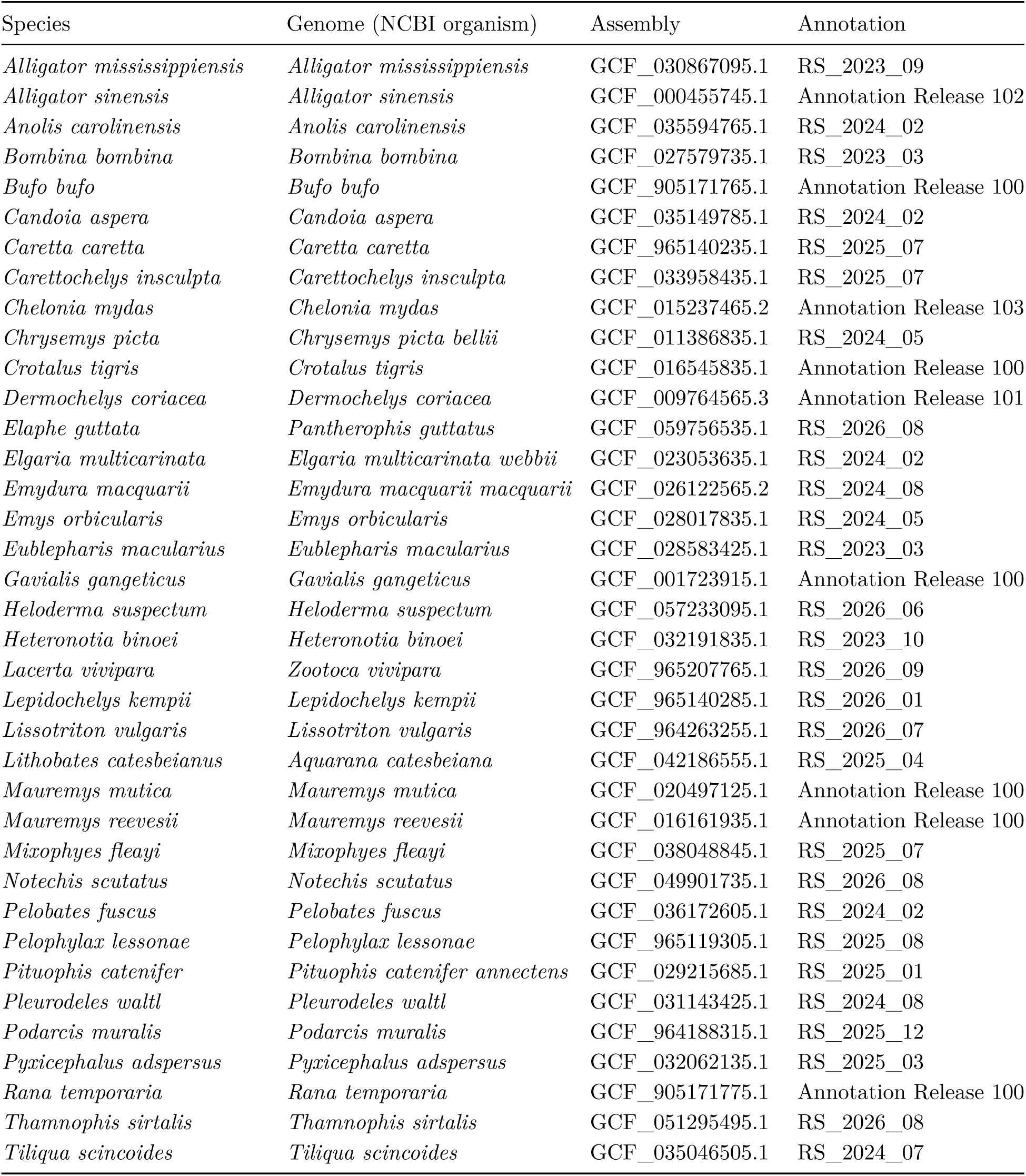
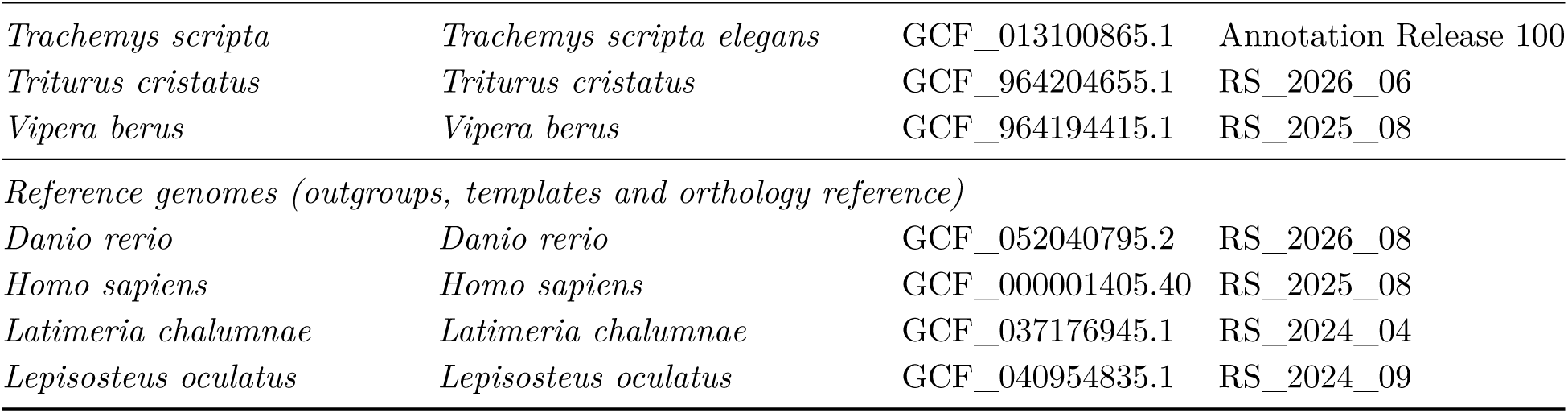
Study 2 genome assemblies and RefSeq annotations (NCBI Datasets). Species are named as in AnAge and Table S7, followed by the NCBI organism of the genome used. The reference genomes are the three fishes used as outgroups and templates and the human genome used as the orthology reference and as an alternative template.

## References

[1] Beth A. Reinke, Hugo Cayuela, Fredric J. Janzen, et al. Diverse aging rates in ectothermic tetrapods provide insights for the evolution of aging and longevity. Science, 376:1459–1466, 2022. doi: 10.1126/science.abm0151.

[2] Rita da Silva et al. Slow and negligible senescence among testudines challenges evolutionary theories of senescence. Science, 376:1466–1470, 2022. doi: 10.1126/science.abl7811.

[3] Viviana I. Pérez, Rochelle Buffenstein, Venkata Masamsetti, Shanique Leonard, Adam B. Salmon, James Mele, Blazej Andziak, Ting Yang, Yael Edrey, Bertrand Friguet, Walter Ward, Arlan Richardson, and Asish Chaudhuri. Protein stability and resistance to oxidative stress are determinants of longevity in the longest-living rodent, the naked mole-rat. Proceedings of the National Academy of Sciences USA, 106(9):3059–3064, 2009. doi: 10.1073/pnas.0809620106.

[4] Xiao Tian, Denis Firsanov, Zhihui Zhang, et al. SIRT6 is responsible for more efficient DNA double-strand break repair in long-lived species. Cell, 177(3):622–638, 2019. doi: 10.1016/j.cell.2019.03.043.

[5] Xavier Farré, Ruben Molina, Fabio Barteri, Paul R. H. J. Timmers, Peter K. Joshi, Baldomero Oliva, Sandra Acosta, Borja Esteve-Altava, Arcadi Navarro, and Gerard Muntané. Comparative analysis of mammal genomes unveils key genomic variability for human life span. Molecular Biology and Evolution, 38(11):4948–4961, 2021. doi: 10.1093/molbev/msab219.

[6] Tomoko Ohta. Slightly deleterious mutant substitutions in evolution. Nature, 246:96–98, 1973. doi: 10.1038/246096a0.

[7] Konstantin Popadin, Leonard V. Polishchuk, Leila Mamirova, et al. Accumulation of slightly deleterious mutations in mitochondrial protein-coding genes of large versus small mammals. Proceedings of the National Academy of Sciences USA, 104(33):13390–13395, 2007. doi: 10.1073/pnas.0701256104.

[8] Gwen R. Buel and Kylie J. Walters. Can AlphaFold2 predict the impact of missense mutations on structure? Nature Structural & Molecular Biology, 29(1):1–2, 2022. doi: 10.1038/s41594-021-00714-2.

[9] Marina A. Pak, Karina A. Markhieva, Mariia S. Novikova, Dmitry S. Petrov, Ilya S. Vorobyev, Ekaterina S. Maksimova, Fyodor A. Kondrashov, and Dmitry N. Ivankov. Using AlphaFold to predict the impact of single mutations on protein stability and function. PLoS ONE, 18(3): e0282689, 2023. doi: 10.1371/journal.pone.0282689.

[10] Henry Dieckhaus, Michael Brocidiacono, Nicholas Z. Randolph, and Brian Kuhlman. Transfer learning to leverage larger datasets for improved prediction of protein stability changes. Proceedings of the National Academy of Sciences USA, 121(6):e2314853121, 2024. doi: 10.1073/pnas.2314853121.

[11] Kotaro Tsuboyama et al. Mega-scale experimental analysis of protein folding stability in biology and design. Nature, 620:434–444, 2023. doi: 10.1038/s41586-023-06328-6.

[12] João Pedro de Magalhães, Zoya Abidi, Gabriel Arantes dos Santos, et al. Human ageing genomic resources: updates on key databases in ageing research. Nucleic Acids Research, 52 (D1):D900–D908, 2024. doi: 10.1093/nar/gkad927.

[13] João Pedro de Magalhães, Joana Costa, and George M. Church. An analysis of the relationship between metabolism, developmental schedules, and longevity using phylogenetic independent contrasts. Journals of Gerontology Series A: Biological Sciences and Medical Sciences, 62(2): 149–160, 2007. doi: 10.1093/gerona/62.2.149.

[14] Sarah C. Dyer, Olanrewaju Austine-Orimoloye, Andrey G. Azov, et al. Ensembl 2025. Nucleic Acids Research, 53(D1):D948–D957, 2025. doi: 10.1093/nar/gkae1071.

[15] Nathan S. Upham, Jacob A. Esselstyn, and Walter Jetz. Inferring the mammal tree: species-level sets of phylogenies for questions in ecology, evolution, and conservation. PLoS Biology, 17 (12):e3000494, 2019. doi: 10.1371/journal.pbio.3000494.

[16] Nuala A. O’Leary, Eric Cox, J. Bradley Holmes, et al. Exploring and retrieving sequence and metadata for species across the tree of life with NCBI datasets. Scientific Data, 11:732, 2024. doi: 10.1038/s41597-024-03571-y.

[17] Brunno Freire Oliveira, Vinícius Avelar São-Pedro, Georgina Santos-Barrera, Caterina Penone, and Gabriel C. Costa. AmphiBIO, a global database for amphibian ecological traits. Scientific Data, 4:170123, 2017. doi: 10.1038/sdata.2017.123.

[18] Oleksandra Oskyrko, Chunrong Mi, Shai Meiri, and Weiguo Du. ReptTraits: a comprehensive dataset of ecological traits in reptiles. Scientific Data, 11:243, 2024. doi: 10.1038/s41597-024-03079-5.

[19] Sudhir Kumar et al. TimeTree 5: an expanded resource for species divergence times. Molecular Biology and Evolution, 39(8):msac174, 2022. doi: 10.1093/molbev/msac174.

[20] John Jumper, Richard Evans, Alexander Pritzel, et al. Highly accurate protein structure prediction with AlphaFold. Nature, 596:583–589, 2021. doi: 10.1038/s41586-021-03819-2.

[21] Mihaly Varadi, Damian Bertoni, Paulyna Magana, et al. AlphaFold protein structure database in 2024: providing structure coverage for over 214 million protein sequences. Nucleic Acids Research, 52(D1):D368–D375, 2024. doi: 10.1093/nar/gkad1011.

[22] Kazutaka Katoh and Daron M. Standley. MAFFT multiple sequence alignment software version 7: improvements in performance and usability. Molecular Biology and Evolution, 30(4):772–780, 2013. doi: 10.1093/molbev/mst010.

[23] Bui Quang Minh, Heiko A. Schmidt, Olga Chernomor, Dominik Schrempf, Michael D. Woodhams, Arndt von Haeseler, and Robert Lanfear. IQ-TREE 2: new models and efficient methods for phylogenetic inference in the genomic era. Molecular Biology and Evolution, 37(5):1530–1534, 2020. doi: 10.1093/molbev/msaa015.

[24] Si Quang Le and Olivier Gascuel. An improved general amino acid replacement matrix. Molecular Biology and Evolution, 25(7):1307–1320, 2008. doi: 10.1093/molbev/msn067.

[25] Henry Dieckhaus and Brian Kuhlman. Protein stability models fail to capture epistatic interactions of double point mutations. Protein Science, 34(1):e70003, 2025. doi: 10.1002/pro.70003.

[26] Mark Pagel. Inferring the historical patterns of biological evolution. Nature, 401:877–884, 1999. doi: 10.1038/44766.

[27] Robert P. Freckleton, Paul H. Harvey, and Mark Pagel. Phylogenetic analysis and comparative data: a test and review of evidence. The American Naturalist, 160(6):712–726, 2002. doi: 10.1086/343873.

[28] Yoav Benjamini and Yosef Hochberg. Controlling the false discovery rate: a practical and powerful approach to multiple testing. Journal of the Royal Statistical Society: Series B, 57(1): 289–300, 1995. doi: 10.1111/j.2517-6161.1995.tb02031.x.

[29] Benoît Nabholz, Sylvain Glémin, and Nicolas Galtier. Strong variations of mitochondrial mutation rate across mammals—the longevity hypothesis. Molecular Biology and Evolution, 25 (1):120–130, 2008. doi: 10.1093/molbev/msm248.

[30] H. Bradley Shaffer et al. The western painted turtle genome, a model for the evolution of extreme physiological adaptations in a slowly evolving lineage. Genome Biology, 14(3):R28, 2013. doi: 10.1186/gb-2013-14-3-r28.

[31] Richard E. Green et al. Three crocodilian genomes reveal ancestral patterns of evolution among archosaurs. Science, 346:1254449, 2014. doi: 10.1126/science.1254449.

[32] Cuihong Wan, Blake Borgeson, Sadhna Phanse, et al. Panorama of ancient metazoan macromolecular complexes. Nature, 525(7569):339–344, 2015. doi: 10.1038/nature14877.

[33] Anna C. F. Lewis, Nick S. Jones, Mason A. Porter, and Charlotte M. Deane. What evidence is there for the homology of protein-protein interactions? PLoS Computational Biology, 8(9): e1002645, 2012. doi: 10.1371/journal.pcbi.1002645.

[34] Chi-Wei Yeh, Kuan-Lun Hsu, Shu-Ting Lin, et al. Altered assembly paths mitigate interference among paralogous complexes. Nature Communications, 15(1):7169, 2024. doi: 10.1038/s41467-024-51286-w.

[35] Charles R. Harris, K. Jarrod Millman, Stéfan J. van der Walt, et al. Array programming with NumPy. Nature, 585(7825):357–362, 2020. doi: 10.1038/s41586-020-2649-2.

[36] Pauli Virtanen, Ralf Gommers, Travis E. Oliphant, et al. SciPy 1.0: fundamental algorithms for scientific computing in Python. Nature Methods, 17(3):261–272, 2020. doi: 10.1038/s41592-019-0686-2.

[37] Wes McKinney. Data structures for statistical computing in Python. In Proceedings of the 9th Python in Science Conference, pages 56–61, 2010. doi: 10.25080/Majora-92bf1922-00a.

[38] Skipper Seabold and Josef Perktold. statsmodels: econometric and statistical modeling with Python. In Proceedings of the 9th Python in Science Conference, pages 92–96, 2010. doi: 10.25080/Majora-92bf1922-011.

[39] Peter J. A. Cock et al. Biopython: freely available Python tools for computational molecular biology and bioinformatics. Bioinformatics, 25(11):1422–1423, 2009. doi: 10.1093/bioinformatics/btp163.

[40] Adam Paszke, Sam Gross, Francisco Massa, et al. PyTorch: an imperative style, high-performance deep learning library. In Advances in Neural Information Processing Systems 32, pages 8024–8035, 2019.

[41] John D. Hunter. Matplotlib: a 2D graphics environment. Computing in Science & Engineering, 9(3):90–95, 2007. doi: 10.1109/MCSE.2007.55.

[42] J. Dauparas, I. Anishchenko, N. Bennett, et al. Robust deep learning-based protein sequence design using ProteinMPNN. Science, 378(6615):49–56, 2022. doi: 10.1126/science.add2187.

[43] A. Shrake and J. A. Rupley. Environment and exposure to solvent of protein atoms. Lysozyme and insulin. Journal of Molecular Biology, 79(2):351–371, 1973. doi: 10.1016/0022-2836(73)90011-9.

[44] Matthew Z. Tien, Austin G. Meyer, Dariya K. Sydykova, Stephanie J. Spielman, and Claus O. Wilke. Maximum allowed solvent accessibilites of residues in proteins. PLoS ONE, 8(11):e80635, 2013. doi: 10.1371/journal.pone.0080635.

